# A quantitative ultrastructural timeline of nuclear autophagy reveals a role for dynamin-like protein 1 at the nuclear envelope

**DOI:** 10.1101/2024.02.14.580336

**Authors:** Philip J. Mannino, Andrew Perun, Ivan V. Surovtsev, Nicholas R. Ader, Lin Shao, Elisa C. Rodriguez, Thomas J. Melia, Megan C. King, C. Patrick Lusk

## Abstract

Autophagic mechanisms that maintain nuclear envelope homeostasis are bulwarks to aging and disease. By leveraging 4D lattice light sheet microscopy and correlative light and electron tomography, we define a quantitative and ultrastructural timeline of nuclear macroautophagy (nucleophagy) in yeast. Nucleophagy begins with a rapid accumulation of the selective autophagy receptor Atg39 at the nuclear envelope and finishes in ∼300 seconds with Atg39-cargo delivery to the vacuole. Although there are several routes to the vacuole, at least one pathway incorporates two consecutive membrane fission steps: inner nuclear membrane (INM) fission to generate an INM-derived vesicle in the perinuclear space and outer nuclear membrane (ONM) fission to liberate a double membraned vesicle to the cytosol. ONM fission occurs independently of phagophore engagement and instead relies surprisingly on dynamin like 1 (Dnm1), which is recruited to sites of Atg39 accumulation by Atg11. Loss of Dnm1 compromises nucleophagic flux by stalling nucleophagy after INM fission. Our findings reveal how nuclear and INM cargo are removed from an intact nucleus without compromising its integrity, achieved in part by a non-canonical role for Dnm1 in nuclear envelope remodeling.

## Introduction

The nuclear envelope (NE) establishes the boundary between cytoplasm and nucleoplasm. Although it shares a contiguous lumen with the endoplasmic reticulum (ER), it is functionally specialized by the unique proteomes and lipidomes that distinguish at least three membranes: the inner nuclear membrane (INM) that engages with the genome and in some eukaryotes, a nuclear lamina, an outer nuclear membrane (ONM) that connects with the cytoskeleton, and a pore membrane that houses nuclear pore complexes (NPCs). As for all organelles, NE proteins are subject to proteostasis mechanisms that include the ubiquitin-proteasome and autophagy pathways^1^. These degradative processes are triggered upon molecular damage or genetic perturbation to NE components^2–4^, nutrient deprivation^5^, mislocalization of integral membrane proteins to the INM^6–8^, senescence^9^, during the meiotic program^10,11^, and in the context of other cellular stresses^12^. Although turnover of the NE is important to maintain cellular homeostasis by combatting the accumulation of deleterious factors associated with disease and aging^13^, the underlying mechanisms remain to be fully defined.

Of the two major protein degradation pathways, autophagy recycles a broader spectrum of molecules including protein aggregates, nucleic acids, and lipids^14^. The latter may be particularly important at the nucleus as the NE is sensitive to perturbations in lipid composition^15–18^ and metabolism^19–21^. Autophagy delivers molecules (cargo) for degradation in lysosomes (in metazoans) or vacuoles (in yeast and plants) either directly through microautophagy or indirectly through macroautophagy, which incorporates a double membrane autophagosome intermediate^22^. The precursor to the autophagosome, the phagophore or isolation membrane, is a cup-shaped lipid bilayer that forms de novo and expands around cargo with the aid of scaffold proteins like Atg11 in yeast or FIP200 in mammals^23^. There are both non-selective and selective macroautophagy pathways with the latter defined by selective autophagy receptors (SARs) that are recruited to specific cargo and interface with the Atg11/FIP200 scaffolds and membrane-associated proteins on the phagophore, most commonly lipidated LC3 (Atg8 in yeast)^22^.

Both micro and macroautophagy pathways act on the nucleus. For example, mammalian cells execute a form of nuclear microautophagy whereby the ONM selectively buds into lysosomes^24^. Similarly, piecemeal microautophagy of the nucleus (PMN) in yeast involves the coordinated evagination of both the INM and ONM directly into the vacuole^25^. The forms of nuclear macroautophagy described to date are comparatively less well understood and atypical as selectivity is imparted by direct interactions between Atg8/LC3 and the cargo instead of through a dedicated SAR. These pathways include the clearance of Lamin B1 under conditions of oncogene-induced senescence^9^ and NPCs by “NPC-phagy”^26–28^. In contrast, work in budding yeast identified the first nuclear-specific SAR, Atg39^29^, that is required for the degradation of both soluble nuclear and integral membrane cargo^29–31^. However, in this pathway, and indeed all forms of nuclear autophagy, the NE remodeling mechanisms required to selectively remove components of the nucleus, INM, and ONM remain mechanistically opaque.

Because Atg39 is a SAR and is consumed by autophagy^29^, it provides an ideal molecular handle to investigate the kinetics and NE remodeling steps of a model nuclear macroautophagy (hereafter called nucleophagy) pathway. Atg39 is a single pass integral ONM protein^32,33^ with Atg8 and Atg11 interacting motifs on its cytosolic N-terminus^29^. The C-terminal lumenal domain has several amphipathic helices that engage the lumenal leaflet of the INM^32,33^. Thus, Atg39 forms a translumenal bridge that physically couples the INM and ONM and likely coordinates their remodeling during nucleophagy. Consistent with this, the Atg39 lumenal amphipathic helices are required for nucleophagic degradation of nuclear and INM cargos^32,33^. Further, Atg39 overexpression drives elaborate NE blebbing where both membranes of the NE are evaginated^32,33^. However, as these NE blebs remain associated with the NE in the absence of autophagy induction, there remains uncertainty as to the precise membrane remodeling events orchestrated by Atg39 in the physiological state, which must be understood at the ultrastructural and molecular levels.

Two hypothetical models have been proposed for Atg39-mediated nucleophagy^1^. In the first model, two sequential membrane fission events are invoked, first at the INM to generate an INM-derived vesicle (INMDV) in the perinuclear space/NE lumen and the second at the ONM to release a nucleus-derived vesicle (NDV)(Figure S1A). In the second model, the INM and ONM are simultaneously evaginated and an NDV is liberated from the NE through coincident ONM and INM fission events (Figure S1B). Here, we use a combination of lattice light sheet microscopy (LLSM) and correlative light and electron microscopy (CLEM) and tomography to describe the kinetic and ultrastructural steps of nucleophagy. Our data firmly establish that nucleophagy proceeds through an INMDV intermediate. In at least a third of nucleophagy events, INM fission is followed by an ONM fission step that requires Dnm1, a dynamin-like protein that was formerly established to function only in mitochondrial^34^ and peroxisomal^35^ fission. Thus, we significantly advance a molecular and morphological understanding of a nucleophagy pathway.

## Results

### Dnm1 is required for robust nucleophagic flux

To identify membrane remodeling factors that contribute to nucleophagy, we assessed the autophagic flux of Atg39 by modeling an approach used to monitor ER-phagy^36^. We generated yeast strains producing an Atg39 fusion protein with a variation of the Rosella^37^ biosensor that is comprised of a tandem pH-sensitive GFP (pHluorin/pHn) and pH-insensitive mCherry/mCh (Figure 1A, B). Atg39-pHn-mCh was expressed from the *ATG39* chromosomal locus with the endogenous 3’UTR intact. We determined that preserving the *ATG39* 3’UTR was essential for optimal expression of Atg39 fusion proteins, particularly when stimulated under conditions of nitrogen starvation but not Tor1-inhibition using rapamycin (Figure S1C-E).

**Figure 1.**
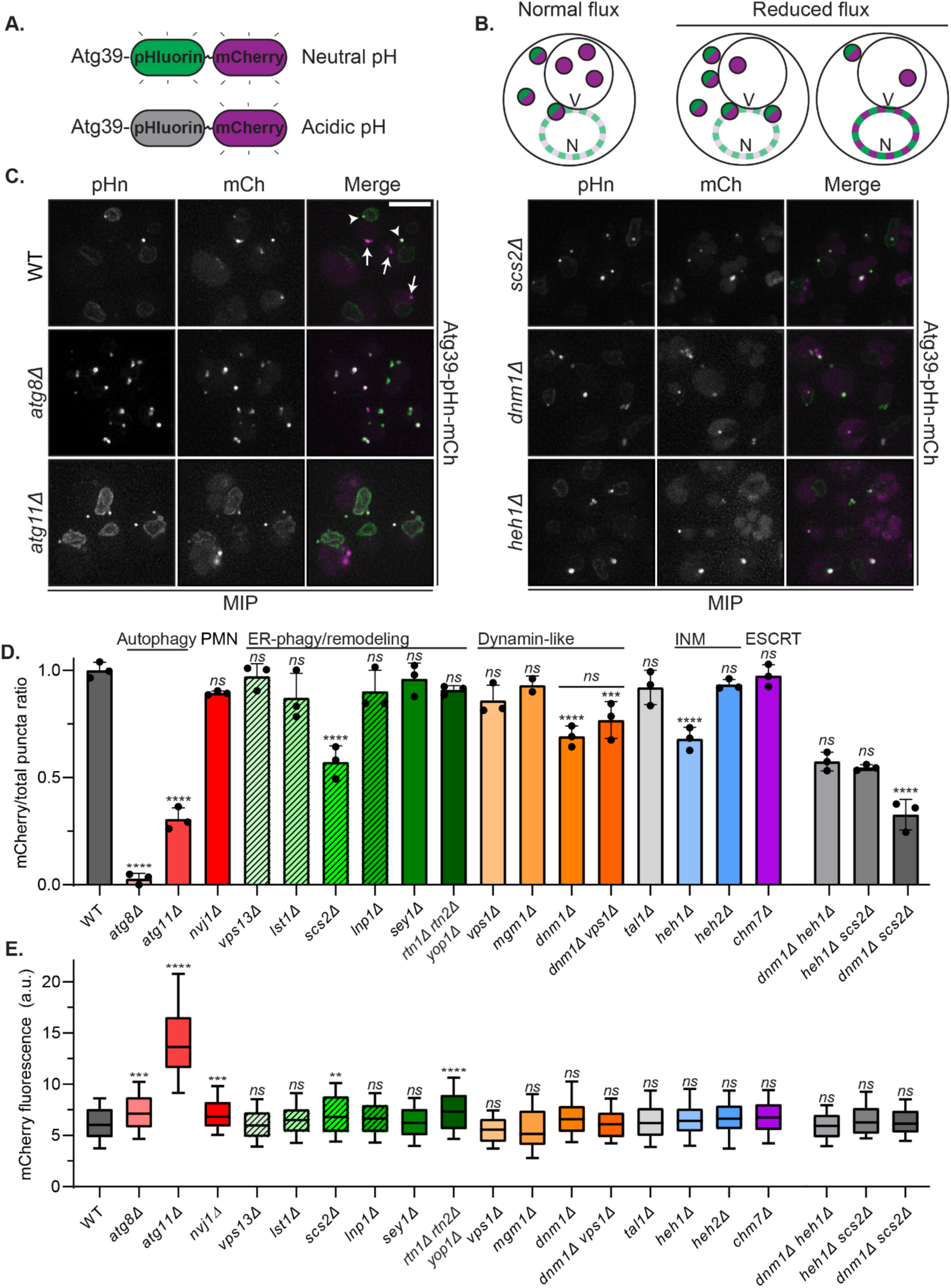
– Dnm1 is required for nucleophagic flux of an Atg39 reporter. **A)** Schematic of Atg39-pHn-mCh construct that has red and green fluorescence in neutral pH and only red fluorescence (magenta) in acidic environments like the vacuole. **B)** Potential outcomes and interpretation of Atg39-pHn-mCh flux experiment. Schematic of cells (N, nucleus; V, vacuole) with Atg39-pHn-mCh foci diagrammed as either green/magenta or magenta; reduced flux can also manifest as an increase in Atg39-pHn-mCh fluorescence at the NE (far right). **C)** Fluorescence micrographs of cells expressing Atg39-pHn-mCh in the indicated genetic backgrounds after 6 h in medium lacking nitrogen. Green (pHn), red (mCh, magenta) channels shown with merge. In all panels a maximum intensity projection (MIP) of a deconvolved z-series of images of entire cell volumes are shown. Arrowheads indicate green/red foci; arrows denote red only foci. Scale bar is 5 µm. **D)** Bar graph of the mean ratios of mCherry-only fluorescent Atg39-pHn-mCh foci to the total number of foci in individual cells of indicated genotypes relative to this ratio in WT cells. Genotype labels are organized by established function (described above the graph) of the proteins they are missing. Dashed lines on the green bar graphs indicate genes required for ER-phagy. Error bars are standard deviation from the mean of three biological replicates (dots) except for *mgm1Δ* cells with two. Total n values for each genotype: WT, 279; *atg8Δ*, 240; *atg11Δ,* 414; *nvj1Δ*, 437; *vps13Δ*, 348; *ist1Δ*, 388; *scs2Δ,* 430; *lnp1Δ*, 374; *sey1Δ*, 331; *rtn1Δrtn2Δyop1Δ,* 371; *vps1Δ,* 349; *mgm1Δ*, *205; dnm1Δ,* 372; dnm1*Δ* vps1*Δ,* 355*; tal1Δ 352; heh1Δ*, 486; *heh2Δ*, 357; *chm7Δ*, *501; dnm1Δ heh1Δ, 355; heh1Δ scs2Δ, 339; dnm1Δ scs2Δ, 335.* Ordinary one-way ANOVA with multiple comparisons with \*\*\*\**P*<0.0001; \*\*\**P*<0.001; *ns*, not significant when either compared to the WT value or, for *dnm1Δheh1Δ* and *dnm1Δscs2Δ* to *dnm1Δ*, or, *scs2Δheh1Δ* to *heh1Δ*. **E)** Box plot of the mCherry fluorescence intensity (in arbitrary units, a.u.) at the nuclear envelope at regions devoid of Atg39-pHn-mCh foci. Top and bottom of the box are the 75^th^ and 25^th^ percentiles, respectively with median indicated by horizontal bar. Error bars indicate the 10-90^th^ percentiles. Data are from three biological replicates except for the *mgm1Δ* cells where data are from two biological replicates. Total n values for each genotype: WT, 150; *atg8Δ*, 150; *atg11Δ,* 151; *nvj1Δ*, 150; *vps13Δ*, 150; *ist1Δ*, 140; *scs2Δ,* 180; *lnp1Δ,* 150; *sey1Δ,* 150; *rtn1Δrtn2Δyop1Δ* cells, 150; *vps1Δ*, 111; *mgm1Δ*, 115; *dnm1Δ*, 135; *dnm1Δvps1Δ, 150; tal1Δ, 150; heh1Δ*, 151; *heh2Δ*, 128; *chm7Δ*, 150; *dnm1Δheh1Δ,* 150*; heh1Δscs2Δ,* 150*; dnm1Δscs2Δ,* 150. Ordinary one-way ANOVA with multiple comparisons. \*\*\*\**P*<0.0001; \*\*\**P*<0.001; \*\**P*<0.01; *ns*, not significant.

After culturing cells without nitrogen, both red-only and red-green Atg39-pHn-mCh fluorescent foci are observed in wildtype (WT) cells (Figure 1C). We calculated a nucleophagic flux value as a ratio of the number of red fluorescent-only foci (i.e. those delivered to the vacuole where the pHn fluorescence is quenched) to the total number of foci in a cell; this ratio was then normalized to 1 for WT cells. We then evaluated nucleophagic flux values in a candidate collection of yeast strains lacking genes that encode nucleophagy cargos (*TAL1, HEH1*)^29,32,33^, contribute to macroautophagy (*ATG8*, *ATG11*)^38,39^, PMN (*NVJ1*)^40^, NE homeostasis (*HEH1*, *HEH2*, *CHM7*)^41–44^, ER-phagy (*VPS13, LST1, SCS2, LNP1*)^45–48^, ER shaping/remodeling (*SEY1, RTN1, RTN2, YOP1*)^49,50^, and membrane fission and fusion mechanisms at other organelles (*DNM1, MGM1, VPS1*)^51–53^. Consistent with the conclusion that this approach reported on perturbations to nucleophagy, Atg39-pHn-mCh flux was nearly abolished in *atg8Δ* cells and severely attenuated in the absence of *ATG11* (Figure 1C, D). In the latter case, we also observed an increase of Atg39-pHn-mCh fluorescence throughout the entire nuclear periphery, suggesting that Atg11 may contribute to early nucleophagy events required for Atg39’s focal accumulation at the NE (Figure 1C, E).

We did not observe any major impact on Atg39-pHn-mCh flux or NE accumulation after perturbing PMN by deletion of *NVJ1* (Figure 1D, E). Likewise, most tested membrane remodeling deletion strains had normal nucleophagic flux, although we note that there were modest increases in NE fluorescence that met reasonable statistical significance in the context of the *rtn1Δrtn2Δyop1Δ* strain (Figure 1E), which has perturbed ER morphology^50^. In marked contrast, we observed a robust 30-45% reduction in nucleophagy in *scs2Δ*, *dnm1Δ*, and *heh1Δ* strains (Figure 1C, D). Of these, Dnm1 (orthologue of DRP1) stood out as it is part of the dynamin family of GTPases that drive membrane fission^54^. Importantly, although Vps1 acts redundantly with Dnm1 in the degradation of peroxisomes^55^, deletion of *VPS1* did not further reduce nucleophagic flux of *dnm1Δ* cells (Figure 1D). While it is known that Dnm1 functions at mitochondria and peroxisomes, there is no prior evidence that it acts at the NE.

We next tested for epistasis between *DNM1*, *HEH1* and *SCS2* by assessing Atg39-pHn-mCh flux in *dnm1Δheh1Δ*, *dnm1Δscs2Δ,* and *scs2Δheh1Δ* strains. As shown in Figure 1D, only the combined deletion of *DNM1* and *SCS2* led to a synergistic loss of nucleophagic flux that was reduced to levels comparable to those in the *atg11Δ* strain. Curiously, deletion of *SCS2* and/or *DNM1* and even *ATG11* did not appreciably impact the total levels of free pHn (‘pHn) generated by vacuolar cleavage of Atg39-pHn-mCherry after 24 h of nitrogen starvation, as assessed by western blot (Figure S1F). Thus, the fluorescence-based assay is more sensitive to measuring nucleophagic flux compared to this end-point assay, but there are clearly even Atg11-independent pathways capable of delivering Atg39 to the vacuole. Two of these pathways are genetically distinguished by deletion of *DNM1* or *SCS2*.

### Dnm1 colocalizes with Atg39 during nucleophagy

Because we are most interested in understanding the mechanisms of membrane remodeling at the NE, we focused primarily on the surprising role for Dnm1 in the degradation of Atg39. In a model where Dnm1 functions during nucleophagy, it is expected to engage Atg39. We therefore tested colocalization between endogenously expressed Dnm1-mCherry and Atg39-GFP during LLSM timelapse imaging (10 second intervals) of live cells under nitrogen starvation (Figure 2). To control for spurious colocalization events between the abundant mitochondrial/peroxisomal-localized Dnm1^34^ and Atg39 in WT cells, we also performed this analysis in the absence of Fis1 (Figure S2), which is required to recruit Dnm1 to mitochondria^51^ and peroxisomes^56^. Further, as we suspected that any interaction between Dnm1 and Atg39 would be fleeting, we also tested how expression of a dominant-negative *dnm1-K41A* allele^57^ that produces a GTPase-dead form of Dnm1 that binds, but does not hydrolyze GTP^57^ impacts any colocalization between Dnm1-mCherry and Atg39-GFP (Figure 2; Figure S2).

**Figure 2.**
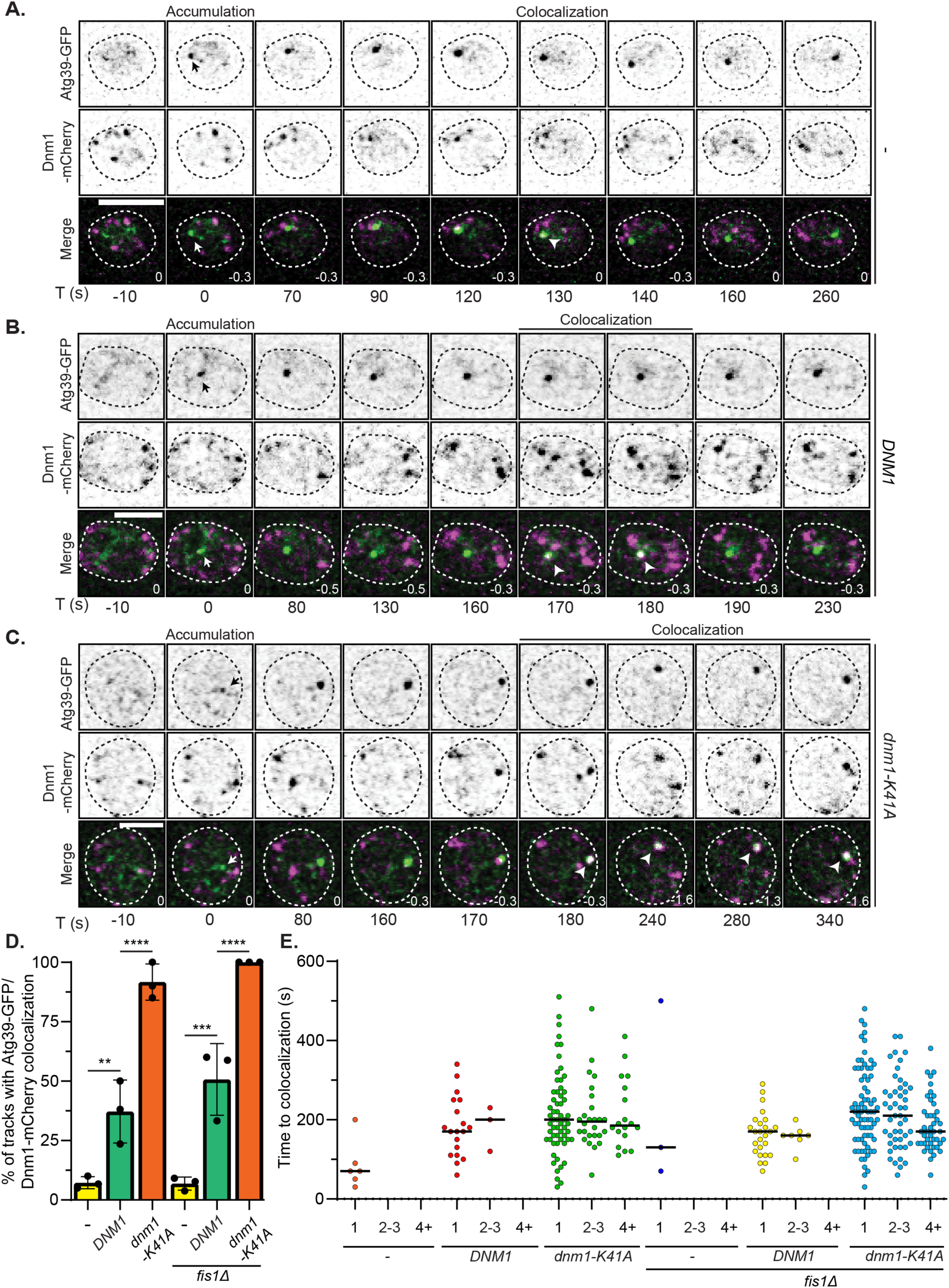
– Dnm1 is recruited to Atg39 during nucleophagy. **A-C)** Inverted fluorescence micrographs of a timelapse series of cells expressing Atg39-GFP and Dnm1-mCherry with and without (-) either an extra copy of *DNM1* or the dominant negative *dnm1-K41A* allele under conditions of nitrogen starvation. Green and red channels shown with merge at indicated times (T) with T=0 s being the first detection of a Atg39-GFP focus (arrow). Arrowheads point to colocalization between Atg39-GFP and Dnm1-mCherry. Numbers in merge are the z-position (in μm) of the image shown in reference to the midplane. Scale bars are 5 µm. **D)** Plot of the percentage of Atg39-GFP foci that were continuously tracked from their point of origin by timelapse microscopy that colocalize with Dnm1-mCherry at least once during a given timelapse image series in either WT or *fis1Δ* cells. Data in each column are presented with mean and standard deviation from 3 biological replicates (dots). Columns 1-6 are *n*=54, 46, 47, 48, 43 and 58 tracks, respectively. Ordinary one-way ANOVA with multiple comparisons. \*\*\*\**P*<0.0001; \*\*\**P*<0.001; \*\**P*<0.01. **E)** A scatter plot of the time between the appearance of an Atg39-GFP focus and colocalization of Atg39-GFP and Dnm1-mCherry. Data are binned by the number of consecutive frames (10 s intervals) colocalization was observed in WT and *fis1Δ* cells with and without expressing *DNM1* or *dnm1-K41A*. The black bar is the median.

We recorded the timing of colocalization relative to the onset of nucleophagy defined as the first observation of an Atg39-GFP focus in a cell. Using this criterion, we observed colocalization between Atg39-GFP and Dnm1-mCherry in less than 10% of WT and *fis1Δ* Atg39-GFP tracks (Figure 2A, D; S2A). Individual colocalization events persisted for only one frame (i.e. less than 20 seconds; Figure 2E). Interestingly, colocalization between Atg39-GFP and Dnm1-mCherry was much more frequent (25-50% of tracks) when we introduced an untagged copy of *DNM1* into the strains (median time of colocalization of 180 s for WT and 160 s for *fis1Δ,* Figure 2B, D; S2B) suggesting that Dnm1-mCherry may not be fully functional in its ability to bind to the NE during nucleophagy without an unaltered counterpart. Not only was colocalization in this context more frequent, but it often persisted for at least 40 seconds (2-3 frames; Figure 2E) suggestive of a sustained physical interaction. Most strikingly, we observed colocalization between Dnm1-mCherry and Atg39-GFP in virtually all cells in the presence of the *dnm1-K41A* allele. Often the colocalization persisted beyond 40 seconds and sometime lasted throughout entire observation window (>350 seconds; median timing of 190 s for WT and 200 s for *fis1Δ,* Figure 2C, E; S2B). These colocalizations persisted even as the pair of foci moved together (Figure 2C; S2C), strongly suggesting that they associated with the same structure. Thus, taken together, there is a robust engagement of Dnm1 with Atg39 during nucleophagy with virtually all sites of Atg39 accumulation being in principle competent to recruit Dnm1.

Consistent with studies demonstrating that Atg11 directly recruits Dnm1 in the context of mitophagy^58^ and pexophagy^55^, we also determined that Atg11 was required for the recruitment of Dnm1 to the NE (Figure S3A, B). Further, we detected specific Bimolecular Fluorescence Complementation (BiFC^59^, Figure S3C) between Atg11 and Dnm1 tagged with either the N or C-termini of the fluorescent protein Venus suggesting a direct physical interaction between these proteins (Figure S3D). Indeed, BiFC signal was lost upon deletion of the last 30 amino acids of Dnm1 which contains the Atg11 binding site^58^ (Figure S3D). Importantly, as BiFC between Dnm1 and Atg11 also occurs at mitochondria and peroxisomes undergoing autophagy^55,58^, we confirmed that a fraction of the BiFC signal was derived from sites of Atg39 accumulation by colocalization with Atg39-ymTurquoise2 (Figure S3D, E). Thus, Atg11 plays a likely direct role in the recruitment of Dnm1 to Atg39 at the NE.

### Quantitative analysis of Atg39 dynamics

To better interpret the timing of Dnm1 recruitment to Atg39, we needed an accounting of the molecular, kinetic, and morphological steps that comprise nucleophagy. We therefore performed LLSM timelapse imaging of Atg39-GFP under conditions of nitrogen starvation in WT cells. To facilitate monitoring the delivery of Atg39-GFP into vacuoles, we performed these experiments in cells expressing Vph1-mCherry to visualize vacuolar membranes. We measured the intensity of Atg39-GFP foci and used commercial (Imaris) and our own particle tracking algorithms^60–62^ to quantitatively evaluate their initial appearance at the NE, their rate of growth, and their delivery to the vacuole – all of which occurred at a rate of 1.4 events/cell/hour (Figure S4D; Movie 1).

Atg39-GFP was first visible diffusely localized along the nuclear rim before it concentrated into a single focus (Figure 3A). Of note, Atg39 always (*n*=114/114) accumulated directly adjacent to the nucleus-vacuole junction (NVJ), suggesting that nucleophagy may occur at a privileged domain on the NE and/or vacuole (Figure 3A). As with other forms of selective autophagy, we observed that mCherry-Atg8 was recruited near-coincidentally to sites of Atg39-GFP NE accumulation (Figure S4A,B). Atg39-GFP fluorescence rapidly increased by over 4-fold and then leveled off by ∼100 seconds before a modest diminishment that likely reflected photobleaching (Figure 3C; blue, circles). A molecular counting strategy based on relating the fluorescence of Atg39-GFP foci to a GFP standard of known copy number (Fta3-GFP^63^)(Figure S1G) indicated that the median number of Atg39-GFP molecules in a given focus at steady state was 134 (Figure 3D; blue) with ∼50-100 molecules added per minute during the initial phase (Figure 3C,D) likely through the coalescence of smaller Atg39-GFP foci that were detected with faster (2 s/frame) imaging (Figure S4F). Interestingly, robust focal accumulation of Atg39-GFP at the NE was dependent on Atg11: we observed a reduction in the number of Atg39-GFP NE foci/hour in *atg11Δ* cells to ∼45% the level of WT or *dnm1Δ* cells (Figure S4C,D). Further, of the Atg39-GFP NE foci that did form, about half had characteristics that were similar to those in WT cells, whereas the other half failed to reach the same apex of fluorescence intensity and ultimately dispersed throughout the NE (“aborted”; Figure S4C,E).

**Figure 3.**
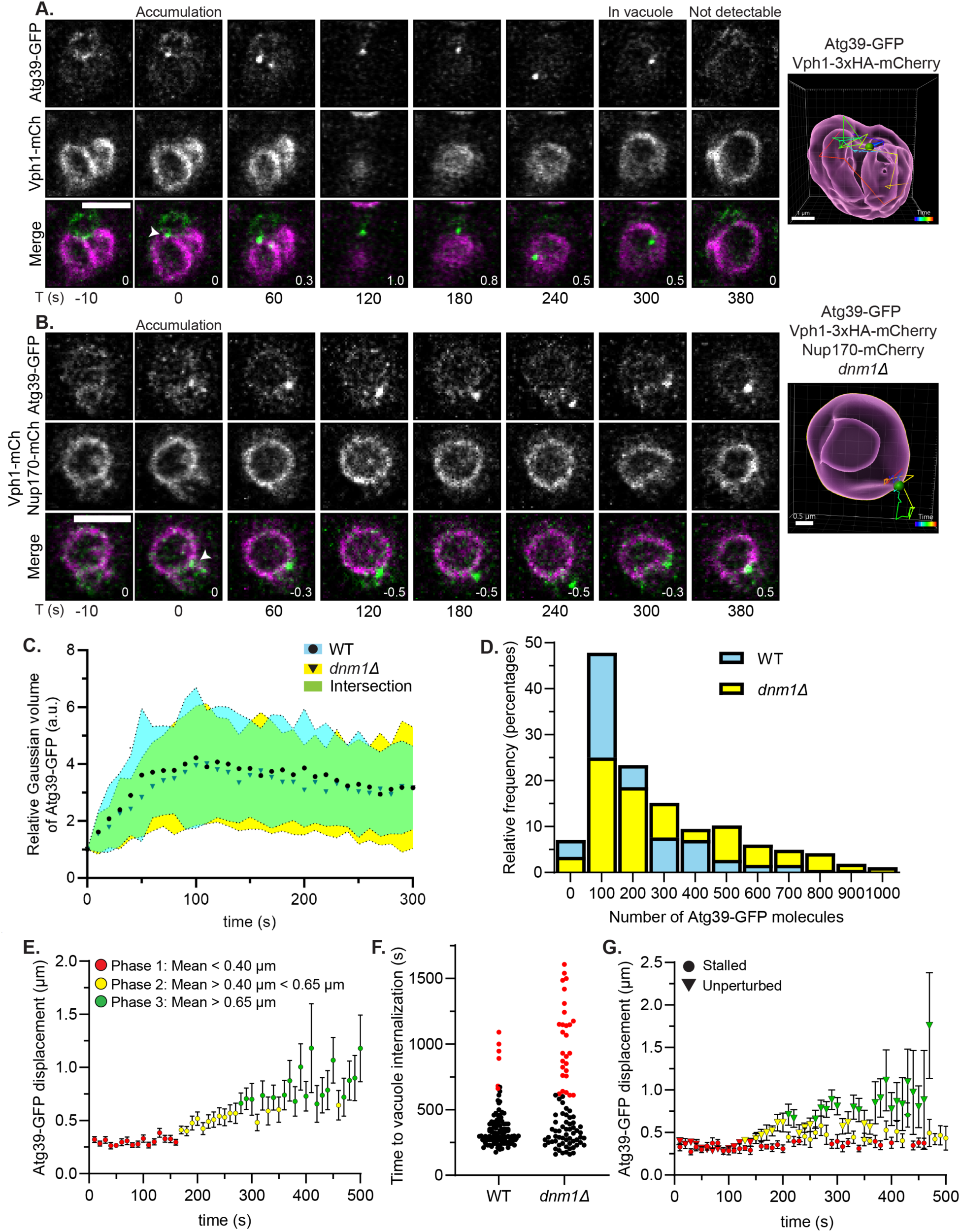
– Dnm1 impacts Atg39 dynamics, copy number, and robust delivery to vacuole. **A)** Fluorescence micrographs from a timelapse series of WT cells expressing Atg39-GFP and Vph1-mCherry (mCh) in media lacking nitrogen. The images shown are at indicated times (T) with T=0 s being the first detection of a Atg39-GFP focus (arrow), which is followed until Atg39-GFP fluorescence disappears in the vacuole (Quenched). Green and red channels shown with merge. Numbers in merge are the z-position (in μm) of the image shown in reference to the midplane. Scale bar is 3 µm. The panel at right shows a 3D model with particle track of Atg39-GFP focus color coded by time as indicated by key. Vph1-mCherry was rendered as a magenta surface and Atg39-GFP as a green ball. Scale bar length is indicated in the image. **B)** As in A but for *dnm1Δ* cells. **C)** Atg39-GFP foci intensity versus time after their first detection. Data are normalized to the volume of the Gaussian of the first detection of an Atg39-GFP focus. Circles (WT) and triangles (*dnm1Δ*) indicate the mean and shaded areas are the standard deviation from three biological replicates. For WT cells, *n*=46 tracks, 45 cells. For *dnm1Δ* cells, *n*=44 tracks, 43 cells. **D)** Histogram of the distribution (% of total) of the number of Atg39-GFP molecules per focus at steady state in cells of the indicated genotypes cultured for 4 h without nitrogen. A total of 184 and 266 Atg39-GFP foci were analyzed from three biological replicates of WT and *dnm1Δ* cells, respectively. **E)** Plot of the mean displacement of Atg39-GFP foci over time in WT cells. Bars are standard error of mean (SEM). The points are colored as indicated in the legend. *n*=40 tracks from 38 cells from three biological replicates. **F)** Scatter plot of the time between the initial identification of an Atg39-GFP focus in WT and *dnm1Δ* cells and internalization into the vacuole. Red dots indicate instances in which the Atg39-GFP particle never entered the vacuole after at least ten minutes. For WT cells, *n*=114 tracks, 99 cells, five biological replicates. For *dnm1Δ* cells, *n*=95 tracks, 82 cells, four biological replicates. **G)** As in **E** but *dnm1Δ* cells. Atg39-GFP tracks were grouped together based on whether they were internalized (unperturbed) or not (stalled) in vacuoles. The points are colored as indicated in the legend in **E.** *n*=37 unperturbed tracks (triangles), 36 cells and *n*=19 stalled tracks (circles), 19 cells, three biological replicates.

To quantitatively describe Atg39’s journey from the NE to the vacuole, we measured the displacements of each of 40 Atg39-GFP foci every 10 seconds over 20-minute timelapse experiments from the first frame of their NE focal accumulation. A consistent theme emerged in which the Atg39-GFP tracks could be empirically categorized into three bins based on characteristic mean displacements for each time point (Figure 3E). In an initial phase that extended until ∼160 seconds, Atg39-GFP foci were constrained with average displacements of 0.30 ± 0.03 µm (Figure 3E; red). This was followed by ∼100 seconds where the average displacement roughly doubled to 0.52 ± 0.08 µm (Figure 3E; yellow). Finally, at ∼280 seconds Atg39-GFP foci entered a highly mobile state with a mean displacement of 0.82 ± 0.18 µm (Figure 3E; green). Thus, Atg39-GFP foci transition from a constrained to a highly mobile state during nucleophagy.

The categorization of Atg39-GFP dynamics into three states provided a useful framework to interpret and temporally align key molecular events. For example, manual inspection of all Atg39-GFP tracks with the vacuole membrane marker (Vph1-mCherry) indicated that delivery into the vacuole lumen occurred within a range of 3 to 8 minutes after Atg39-GFP accumulation with a median of ∼284 seconds (Figure 3A, F), notably like the timing of the transition from phase 2 to 3. Likewise, the transition from phase 1 to 2 occurs with timing that corresponds to the median timing of Dnm1 recruitment (∼170 seconds after Atg39-GFP accumulation, Figure 2E). Further, and consistent with the interpretation that phase 1 reflects the dynamics of Atg39 at the NE, we observed that a model INM cargo (Figure S5A) colocalizes with Atg39-GFP foci with a mean timing of ∼80 seconds after Atg39-GFP accumulation (Figure S5B, C). Thus, together these data predict that Dnm1 contributes to a key transition step in nucleophagy that corresponds to a step after cargo delivery and between the NE and vacuole.

### Nucleophagy stalls in the absence of Dnm1

To explore a model in which Dnm1 acts during nucleophagy, we monitored Atg39-GFP dynamics in *dnm1Δ* cells. Interestingly, early events including the formation of the Atg39-GFP foci proximal to the NVJ (Figure 3B) and the initial rate of Atg39-GFP accumulation (Figure 3C; yellow, triangles) were indistinguishable between WT and *dnm1Δ* cells. However, analyzing Atg39-GFP copy number within individual foci at steady state revealed an increase to a median of 264 molecules in *dnm1Δ* cells compared to 134 in WT cells (Figure 3D; yellow). Taken together, these data suggest that *dnm1Δ* cells contain a population of Atg39-GFP foci that accumulate additional molecules in a timeframe beyond 300 seconds, implying their delivery to the vacuole may be impaired.

Our prior observations suggested only a partial compromise in nucleophagic flux in the absence of *DNM1* (Figure 1D), and indeed, in the instances where Atg39-GFP was delivered to the vacuole in *dnm1Δ* cells, it did so with timing comparable to WT cells (Figure 3F; black dots). However, 33% of Atg39-GFP foci that were tracked from their NE accumulation forward failed to enter the vacuole over the experimental timeframe compared to only 6% in WT cells (Figure 3F; red dots, Figure S5D). Thus, consistent with the nucleophagic flux assay, these observations suggest that Dnm1 is required for the delivery of at least a third of Atg39 to the vacuole. We next assessed the dynamics of Atg39-GFP foci in *dnm1Δ* cells. As our data suggested at least two fates of Atg39 in *dnm1Δ* cells, we measured the displacement of the “stalled” (Dnm1-dependent; Figure 3G; circles) and “unperturbed” (Dnm1-independent; Figure 3G; triangles) Atg39-GFP tracks separately and color-coded the data to reflect the three states defined in Figure 3E. Strikingly, whereas the unperturbed tracks exhibited dynamics that were similar to WT cells, the stalled tracks had mean displacements that mirror those in the first phase, consistent with the notion that the transition from phase 1 to phase 2 was disrupted by loss of *DNM1* in these cells (Figure 3G; circles). Indeed, the relative time that the stalled Atg39 tracks spent in phase 1 doubled compared to both WT cells and the unperturbed tracks in *dnm1Δ* cells (Figure S5E). Thus, these data support a model in which Dnm1 is required to release Atg39 from the confines of the NE in at least a third of all nucleophagy events.

### Nucleophagy proceeds through a lumenal vesicle intermediate

We next sought to correlate Atg39’s dynamics in WT and *dnm1Δ* cells to an ultrastructural description of nucleophagy by CLEM and tomography. This analysis was enabled by the recently developed hyper folder YFP (hfYFP), which retains more fluorescence compared to mEGFP following fixation, staining, and embedding^64^. We co-expressed Nup170-mCherry and Vph1-mCherry to provide additional landmarks to determine the position of Atg39-hfYFP relative to the nucleus and vacuole, respectively. As we lack an approach to synchronize nucleophagy, we acquired tomograms from randomly selected Atg39-hfYFP foci, avoiding those that were within the vacuole lumen. As CLEM is laborious and low-throughput, we used probability theory to establish a target of ∼30 tomograms, which would allow us to capture at least one image representative of all morphological steps that last longer than 30 seconds given that it takes ∼300 seconds to complete nucleophagy (see methods). Thus, morphologies comprising phase 1 (∼160 seconds) and 2 (∼100 seconds) should be highly represented and perhaps those invisible to the quantitative light microscopy analysis could also be revealed.

We obtained 32 tomograms in WT cells where we correlated the fluorescence of Atg39-hfYFP to the underlying ultrastructure (Figure 4A-E; Figure S6A-E). We observed several morphologies that are schematized with categorization labels (1-4) in Figure 4F; the number of tomograms representative of each category is plotted in Figure 4G (blue bars). We made several observations: 1) Approximately half (15/32) of the Atg39-hfYFP foci correlated to regions of NE (Figure 4A) or, surprisingly, ER (Figure 4B; Figure S6A-C), with intralumenal vesicles that we interpret as INMDVs; in the latter we often visualized a nearby ER-NE junction within the 3D volume of the tomogram (e.g. Figure 4B, see 3D model) while some appeared in the cortical ER proximal to the plasma membrane (Figure S6C). The observation of INMDVs confirmed that INM fission occurs prior to ONM fission (Model 1; Figure S1A). Of additional interest, INMDVs at the NE were associated with an expansion of the ONM (Figure 4A) suggesting that these events may occur alongside membrane biosynthesis and/or recruitment mechanisms. 2) There were no structures that resembled a phagophore near any Atg39-hfYFP focus that correlated to INMDVs in the NE or ER (Category 1). Moreover, although 6 tomograms contained apparent NDVs that had been presumably liberated from the NE/ER (Category 2; Figure 4C), only one of these revealed what appeared to be a phagophore in the process of engulfing the structure (Figure S6D). We did, however, observe 4 Atg39-hfYFP foci that correlated to a vesicle enclosed by 4 membranes, which we interpret to be an NDV within an autophagosome (Category 3; Figure 4D). 3) Atg39-hfYFP foci that were observed to be proximal to the vacuole by fluorescence demarked several interesting morphologies that suggest different fates for the autophagosome containing an NDV. For example, we observed close apposition/contact sites between the vacuole and autophagosome (Figure 4D; arrowhead, Movie 2), which was evocative of a kiss and run-type mechanism that might deliver vacuolar degradative enzymes in addition to the more expected direct vacuole fusion (Category 4; Figure 4E). We also observed Atg39-hfYFP foci that overlapped with Vph1-mCherry signal that correlated with a vesicle containing membrane fragments that we presume to be derived from an NDV and autophagosome (Figure S6E). 4) Lastly, we noted that the diameters of the INMDVs were 25-125 nm when found within the NE/ER lumen (with a few exceptions) but were 200-350 nm within NDVs in the cytosol (Figure 4H; blue dots). Making the reasonable assumption that the NDVs arose from the INMDVs in the NE/ER lumen, the implication is that the INMDVs may fuse together before they are released as NDVs into the cytosol. Indeed, we occasionally observed multiple closely apposed INMDVs in the NE/ER lumen that correlated with a single Atg39-hfYFP focus (Figure 4A, Figure S6A) including one example in which we captured two vesicles connected by a narrow membrane neck as if undergoing fusion or fission (Figure S6B), further supporting this idea. If smaller INMDVs fuse before they are liberated within a cytosolic NDV, a prediction would be that larger vesicles would accumulate in the NE/ER lumen upon inhibition of ONM/ER scission.

**Figure 4.**
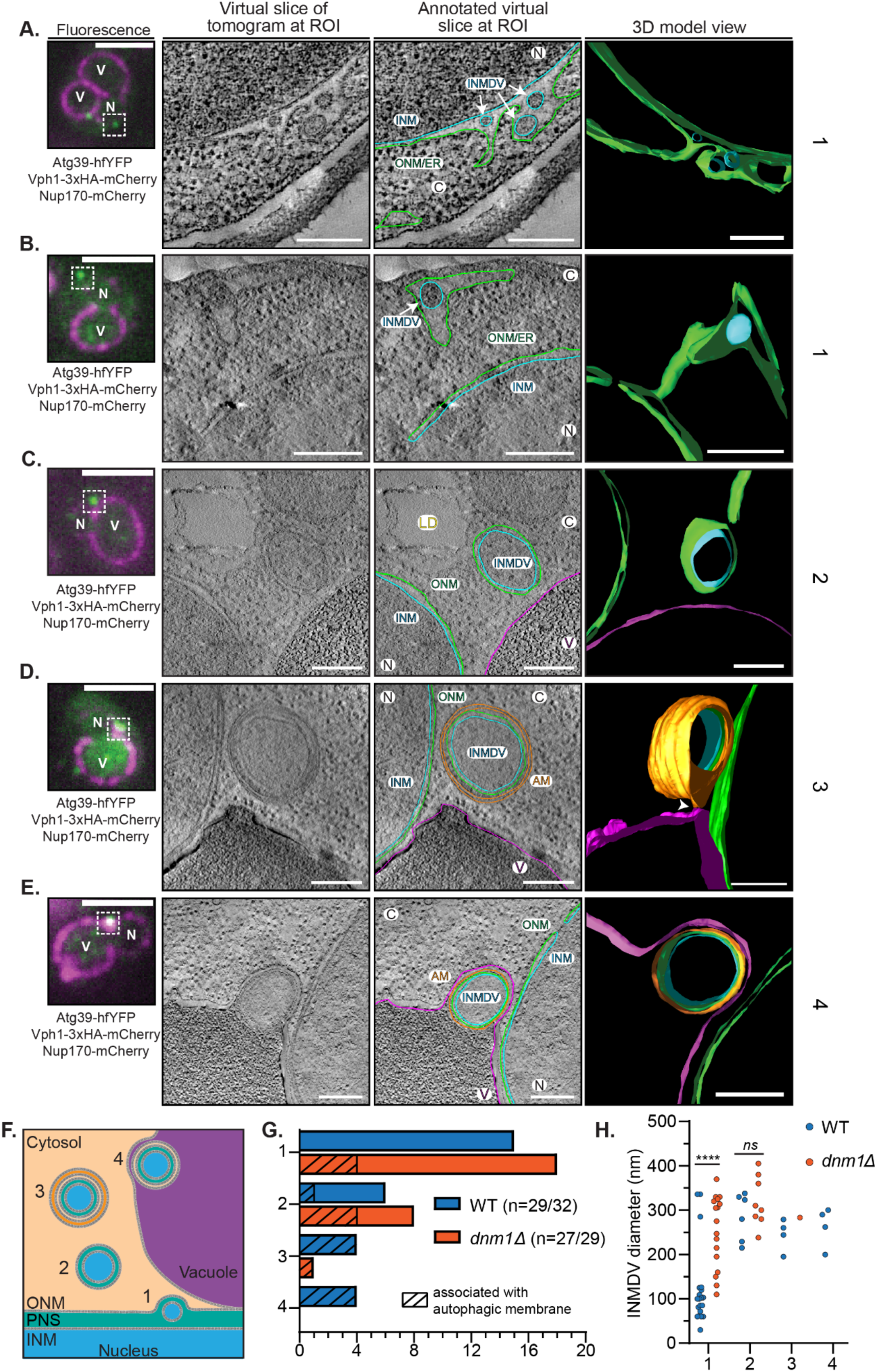
– Nucleophagy proceeds through a lumenal vesicle intermediate. **A-E)** CLEM/tomography of sites of Atg39-hfYFP foci. From left to right: fluorescence image of WT cells expressing Atg39-hfYFP (green), Vph1-mCherry and Nup170-mCherry (magenta). Scale bar is 3 µm. EM tomogram acquired of boxed region in fluorescence image and tomogram virtual slice without and with annotation shown alongside a snapshot of a 3D model. Arrows point to INMDVs. ONM is green, INM is light blue, phagophore/autophagosome is orange, vacuole membrane is magenta. Scale bar is 200 nm. C, cytosol; N, nucleus; V, vacuole; ONM, outer nuclear membrane; INM, inner nuclear membrane; ER, endoplasmic reticulum; INMDV, inner nuclear membrane derived vesicle; AM, autophagic membrane; LD, lipid droplet. Numbers at right refer to classification scheme in **F**. Arrowhead indicates a site of close apposition between the outer membrane of the autophagosome and the vacuole in **D**. **F)** Schematic representation of the morphologies observed at Atg39-hfYFP foci, which are classified by number to facilitate quantification. **G)** Bar graph of the number of tomograms with observed morphology categories (**F**) in WT and *dnm1Δ* cells. Dashed lines indicate the observation of phagophore membranes. Tomograms were acquired at 32 Atg39-hfYFP foci and 29 Atg39-hfYFP foci from multiple thick sections of a WT and *dnm1Δ* embedded sample, respectively. **H)** Scatter plot of INMDV diameters in each morphological category described in **F** in WT and *dnm1Δ* cells. For category 1, *n*=20 vesicles in WT and *n*=17 vesicles in *dnm1Δ* cells. For category 2, *n*=6 vesicles for WT and *n*=8 vesicles for *dnm1Δ* cells. For category 3 *n*=4 vesicles for WT and *n*=1 vesicle for *dnm1Δ* cells. For category 4 *n*=4 vesicles for WT and *n*=0 vesicles for *dnm1Δ* cells. Unpaired two-sided t tests, \*\*\*\**P*<0.0001; *ns*, not significant.

### Dnm1 is required for ONM scission

To better understand how Dnm1 contributes to nucleophagy, we next performed CLEM to localize 29 randomly selected non-vacuolar Atg39-hfYFP foci in the context of the ultrastructure in *dnm1Δ* cells. We categorized the observed morphologies as in WT cells (Figure 4F). Consistent with the idea that nucleophagy is stalled in the absence of *DNM1*, we did not observe any examples of autophagosome-vacuole fusion (Category 4; Figure 4G; red). Instead, there was an increase in the labeling of structures that comprise early events like those containing INMDVs within the NE/ER lumen (Category 1; Figure 4G, Figure 5A, B, Figure S7A, B). Note that this class also incorporates INMDVs found within the cortical ER suggesting that a pathway for INMDVs to travel throughout the ER lumen is retained in the *dnm1Δ* cells (Figure S7E). Most strikingly, unlike in WT cells where we did not observe engagement of phagophores with Atg39-hfYFP positive morphologies that labeled INMDVs in the NE/ER, a quarter of category 1 tomograms had membranes that appeared as ribbons in the 3D segmentation of the whole tomogram volume that were in close apposition to and followed the contour of the ONM (Figure 5C-E, Figure S7C, Movie 3). In several examples as in Figure 5D and 5E, the phagophores surrounded (but failed to fully encompass) the NE-derived membranes. Indeed, the phagophore rim appeared obstructed by NE which would need to be cleared by a membrane fission mechanism for phagophore closure to occur. Consistent with this interpretation, we only observed a single Atg39-hfYFP focus within a fully sealed autophagosome in this dataset (Figure 4G). Interestingly, we also observed a four-fold increase in structures we identified as NDVs surrounded by phagophores in *dnm1Δ* cells compared to WT cells (Figure 4G, Figure S7D), although it is likely that at least some of these structures are category 1 but are miscategorized because their NE/ER connections are outside of the tomogram volumes.

**Figure 5.**
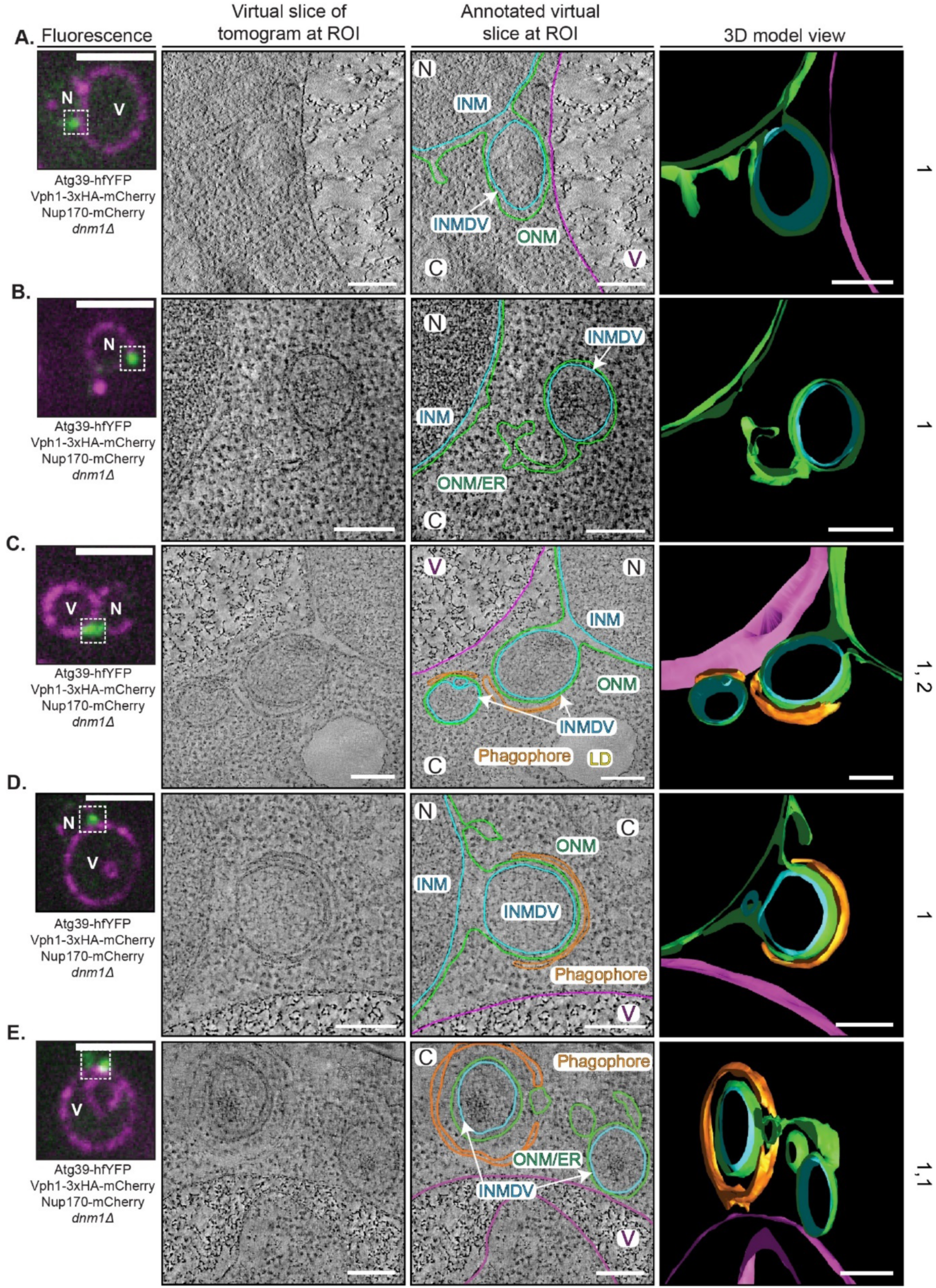
– Dnm1 is required for ONM fission. **A-E)** CLEM/tomography of sites of Atg39-hfYFP foci in *dnm1Δ* cells. From left to right: fluorescence image of *dnm1Δ* cells expressing Atg39-hfYFP (green), Vph1-mCherry and Nup170-mCherry (magenta). Scale bar is 3 µm. EM tomogram acquired of boxed region in fluorescence image and tomogram virtual slice without and with annotation shown alongside a snapshot of a 3D model. Arrows point to INMDVs. ONM/ER is green, INM is light blue, phagophore/autophagosome is orange, vacuole membrane is magenta. Scale bar is 200 nm. C, cytosol; N, nucleus; V, vacuole; ONM, outer nuclear membrane; INM, inner nuclear membrane; ER, endoplasmic reticulum; INMDV, inner nuclear membrane derived vesicle; AM, autophagic membrane. Numbers at right refer to classification scheme in Figure 4F.

Additional evidence for stalling of nucleophagy at a step where membrane fission mechanisms must be invoked is also apparent in the sizes of the INMDVs in *dnm1Δ* cells. Indeed, although in WT cells INMDVs within the NE/ER lumen tended to be 25-125 nm in diameter (Category 1; Figure 4H), they were significantly larger (from 100-400 nm; Figure 4H; red dots) in *dnm1Δ* cells and resembled those that were found only in WT cell NDVs, i.e. after they had been released from the NE/ER. These data thus suggest that a delay in an ONM fission mechanism would allow the buildup of additional membrane and perhaps cargo in the INMDV. Such an idea is also supported by the observation that there is a higher copy number of Atg39-GFP within individual foci in *dnm1Δ* cells (Figure 3D). Thus, we conclude that Dnm1 executes ONM fission.

## Discussion

We have determined the kinetic, molecular, and morphological events that comprise selective Atg39-mediated nucleophagy in budding yeast that is summarized in Figure S8. At the level of the light microscope, nucleophagy begins with rapid (∼50-100 molecules/min) local accumulation of Atg39 at a site adjacent to the NVJ, which likely occurs concomitantly with cargo capture, and is followed by its delivery to the vacuole in approximately 5 minutes (Figure 3F). We estimate, based on the surface area of INM that is captured in the INMDVs and the rate of nucleophagy (1.4/cell/hour; Figure S4D), that ∼80% of the INM is recycled over a 24-hour period when cells are starved of nitrogen. Thus, nucleophagy may help rejuvenate the INM.

The data support that there are at least two parallel nucleophagy pathways responsible for the ultimate delivery of Atg39 to the vacuole, one of which is dependent on Dnm1 and the other on Scs2 and likely other yet to be defined factors. As virtually all CLEM micrographs of Atg39 foci revealed the presence of intralumenal vesicles that we interpret to be INMDVs, we suggest that while both pathways incorporate an INM fission mechanism that likely requires the lumenal amphipathic helices of Atg39^32^, the Dnm1-dependent nucleophagy pathway requires a second membrane fission reaction executed at the ONM that liberates an NDV to the cytosol. Our data strongly implicate Dnm1 as a key player in ONM fission. This conclusion is supported by several data including the robust association of Dnm1 with Atg39-containing structures (Figure 2) and the buildup of likely stalled nucleophagy intermediates at the NE in *dnm1Δ* cells (Figure 3; 5). Interestingly, the data also support that a C-terminal fusion of Dnm1 cannot fully engage the ONM as it does at other organelles like mitochondria^65^. Indeed, we only observe robust association of Dnm1-mCherry with Atg39-GFP in the presence of endogenous Dnm1 or the GTP-locked dominant negative. We speculate that there may be ONM-specific constraints that preclude engagement by the Dnm1-mCherry fusion - perhaps an inability to form a polymer that can encircle an ONM neck. Although the nature of such constraints remains to be discovered, they hint that the ultimate mechanism of ONM fission driven by Dnm1 may have unique characteristics at the NE that await discovery.

Despite the potential for a unique Dnm1-mediated fission mechanism at the NE, the data support that the mechanism of Dnm1 recruitment to Atg39 is analogous to how Dnm1 is recruited to sites of Atg32-mediated mitophagy and requires a biochemical interaction between Dnm1 and Atg11^58^. This conclusion is supported by the requirement of Atg11 for Dnm1 recruitment to the NE and the observation of specific BiFC between Atg11 and Dnm1 (that colocalizes with Atg39) and that depends on an Atg11-interacting motif in Dnm1 (Figure S3C, D). How this interaction is ultimately regulated to catalyze ONM fission is not known but may rely on the coordination between Atg11’s two apparent roles during nucleophagy. Specifically, in addition to recruiting Dnm1, Atg11 contributes to clustering of Atg39 (Figure S4C) to help enforce that Atg39 copy number reaches a critical threshold before it is liberated from the NE, which we suggest is around 150 molecules (Figure 3 C, D). As the copy number is stable after it is released from the nucleus (until its vacuolar degradation), we surmise that Atg11 recruits Dnm1 once this critical threshold copy number of Atg39 is reached. In an alternative but non-mutually exclusive model, if we impose a correlation between Atg39 copy number and INMDV size (with the consideration that INMDVs within the NE/ER lumen are typically small, from 50-100 nm with only a handful at 300 nm (Figure 4H)), a reasonable conclusion is that Dnm1 acts once the INMDVs reach ∼200-300 nm in diameter, well within the experimentally-determined size of cytosolic NDVs (median of 300 nm). Such a conclusion is also supported by the observation that we only detect larger diameter INMDVs stuck at the NE when ONM fission is inhibited (Figure 4H). Thus, future studies will be needed to explore the relationship between Atg39 copy number and/or INMDV size and how these factors influence the Atg11-dependent recruitment of Dnm1 for ONM scission.

That ONM scission may only occur once an INMDV reaches a certain size raises the question of the mechanism of INMDV growth. Indeed, as there appears to be a progression from 50-100 nm to much larger 200-400 nm diameter vesicles in the NE/ER lumen, we must entertain the possibility that these vesicles enlarge by INMDV fusion with the caveat that we cannot fully rule out that we failed to ascertain the fate of the smaller vesicles. Although direct evidence for INMDV fusion is scant, one could interpret the quanta-like merging of Atg39-GFP foci during Atg39 growth within such a framework (Figure S4F). Further, we observed one example of two INMDVs fusing (Figure S6B). As we expect INMDV fusion to be a short-lived event, even capturing this single snapshot speaks to its potential prevalence. How INMDV fusion is achieved is a mystery but would almost certainly require a protein fusogen on the lumenal face of the vesicle membrane. Should such a fusogen exist, one must then consider that INMDVs could also fuse back to the INM or with the ONM or ER membranes. This idea is particularly interesting in the context of so-called NE budding/egress pathways where nuclear cargo (e.g. viruses^66^ or large ribonucleoproteins^67^) are packaged into lumenal vesicles that then fuse with the ONM to allow for their cytosolic delivery. Whether there is a functional relationship between NE budding and nucleophagy will require further study and better molecular markers of the NE budding process in other model systems.

To continue the analogy with NE budding, we also made the surprising observation of INMDVs within the cortical ER in both WT (Figure S6C) and *dnm1Δ* (Figure S7E) cells. As we only observe a stalling of one third of nucleophagy events in *dnm1Δ* and the associated Atg39-GFP foci appear stuck at the NE, we propose that INMDVs that escape the ONM and enter the broader ER lumen cannot be targeted by Dnm1 and must therefore be degraded by other ER-phagy type pathways that would be predicted to depend on Scs2 (Figure S8A) or other Atg11-independent mechanisms that await discovery (Figure 1; Figure S1F). Consistent with a role for Scs2, it has been implicated in ER-phagy of the cortical ER at sites of endocytosis^48^. It will be exciting to understand whether there is a functional decision made by cells to engage the Dnm1 or Scs2-dependent pathways, but it is worth noting that we often observe INDMVs being generated into the lumen of junctions between the ER and NE (Figure 4A)^32^. Thus, it is possible that under these circumstances there is a direct delivery of some INMDVs into the ER lumen that allows them to escape Dnm1-targeting.

That there may be at least two pathways that allow INMDVs to exit the NE, one through Dnm1 and one through NE-ER junctions into the cortical ER, suggests additional mechanistic differences between nucleophagy and other organelle selective autophagy pathways. Indeed, most selective autophagy pathways are thought to occur through a tight coupling of cargo capture by SARS with Atg8-coupled phagophore membranes that is coordinated by scaffold proteins like Atg11^23^. Curiously, however, even though we do observe the near simultaneous recruitment of Atg8 at the point of Atg39 local accumulation at the NE by light microscopy (Figure S4A, B), we failed to detect any obvious membranes that resemble growing phagophores at the NE of WT cells by CLEM. There are several plausible explanations for these potentially contradictory results. First, the small vesicles that may make up nascent phagophores may simply be undetectable in the EM. We do not favor this interpretation as our EM is of high quality with excellent preservation and staining of membranes. Second, Atg8 could be directly conjugated to the ONM (or to proximal vacuole membranes) reminiscent of the proposed conjugation of LC3 to the INM during the autophagic degradation of Lamin B1^9^. Finally, non-lipidated Atg8 could associate with Atg39 or with the preautophagosomal structure, the site of autophagosome assembly, perhaps in a poised state^68^ ready to be rapidly conjugated onto a nascent phagophore after the release of an NDV by Dnm1. In sum, our study provides a compelling explanation for how nuclear and INM cargo can be delivered to vacuoles while maintaining nuclear envelope integrity through an unexpected ONM fission mechanism mediated by Dnm1.

## Acknowledgments

We thank the Center for Cellular and Molecular Imaging for assistance with electron microscopy, particularly M. Graham, Z. Zuo, and X. Liu. We are grateful to C. Kraft, M. Graef and L. Lackner for yeast strains. Thank you to K. Li for technical support. This work was funded by the following grants from National Institutes of Health (NIH): R56AG071201 and R21 AG058033 to C.P.L. and T.J.M.; R01 GM105672 to CPL; F32 GM139285 to N.R.A.; F31 AG069490 to P.J.M.

## Author Contributions

Conceptualization: P.J.M., C.P.L.; Methodology: P.J.M, A.P., I.S., N.R.A., L.S., E.R.; Investigation: P.J.M., A.P., I.S., N.R.A.; Validation: P.J.M., A.P.; Data curation: P.J.M., A.P., I.S., N.R.A.; Formal Analysis: P.J.M., A.P., I.S., N.R.A.; Funding acquisition: P.J.M., T.J.M., C.P.L.; Visualization: P.J.M., A.P.; Project administration: P.J.M., M.C.K., C.P.L.; Resources: L.S., C.P.L.; Software: A.P., I.S., L.S.; Supervision: T.J.M., M.C.K., C.P.L.; Writing – original draft: P.J.M., C.P.L.; Writing – review and editing: All authors

## Materials and Methods

### Yeast culturing conditions and autophagy induction

All experiments were performed at 30°C. Budding yeast (*S. cerevisiae*) were cultured in YPA (1% Bacto-yeast extract (Y), 2% Bacto-peptone (P), 0.025% adenine hemi-sulfate (A; Sigma)) supplemented with 2% glucose (D; Sigma) or in csm (complete supplement mixture; Sunrise Science Products) to maintain selection for centromeric plasmids. For one experiment, we used S. pombe strain MKSP3186, which was cultured overnight in YE5S (yeast extract (YE), 250 mg/L adenine, histidine, leucine, uracil, and lysine hydrochloride).

To induce autophagy by nitrogen starvation, cells in log phase were washed and resuspended in synthetic defined medium lacking nitrogen (sd-N) (0.17% Difco Yeast Nitrogen Base without amino acids and ammonium sulfate, and 2% D). Samples were collected at time points indicated in the figures and were prepared for extraction of RNA, fluorescence microscopy, or immunoblotting as described below.

To induce autophagy by inhibiting Tor1 kinase by treatment with rapamycin, rapamycin (in DMSO; Sigma-Aldrich) was added to cells in log phase to a final concentration of 250 ng/mL. Samples collected at time points indicated in the figure legend were prepared for extraction of RNA for RT-qPCR.

To block the vacuolar degradation of Atg39-pHn-mCh by treatment with PMSF, PMSF (in DMSO) was added to cells in log phase to a final concentration of 1mM.

### Yeast strain construction

All strains used in this study are listed in Supplementary Table 1. PCR-based methods^69^ using the pFA6a^70^ and pK3F^71^ plasmid series as templates were used to generate genomic integration cassettes of sequences encoding fluorescent protein genes and to delete ORFs at the genomic loci^72^. To generate PMCPL576, the *mEGFP* gene followed by a kanamycin resistance cassette (*KAN-MX6*) flanked by loxP sites was amplified from pPM09 using primers with homology arms that are complementary to flanking sequences of the region targeted for genomic insertion. The PCR product was transformed into W303A and kanamycin-resistant colonies were selected. A colony containing the *KAN-MX6* cassette integrated into the proper location as assessed by PCR of genomic DNA was transformed with the centromeric plasmid pSH47^73^ that expresses the Cre recombinase under the control of a galactose-inducible (*GAL1*) promoter. Transformants were selected on csm plates lacking uracil (ura). Overnight cultures of individual colonies grown in csm -ura supplemented with 2% raffinose were diluted into csm -ura supplemented with 2% galactose to induce the expression of the Cre recombinase for 5 hours before single cells were plated on YPD. Colonies were screened for loss of *KAN-MX6* by testing growth on geneticin (Thermo Fisher Scientific)-containing plates; the expression of *ATG39-mEGFP* was confirmed by fluorescence microscopy and immunoblot. PMCPL578, PMCPL1257, PMCPL1045, and PMCPL1097 were generated in a similar manner using pPM26, pPM31, pPM28, and pPM27 respectively as PCR templates.

Genomic integration of *DNM1* and *dnm1 K41A* genes (PMCPL1319, PMCPL1087, PMCPL1112, and PMCPL1113) were generated by genomically integrating linearized pPM11 and pPM15 at the *URA3* locus.

### Plasmid generation

All plasmids used in this study are listed in Supplementary Table 2. All DNA fragments used for cloning were purified using a gel purification kit/protocol (Qiagen). All PCR was performed using KOD polymerase (EMD Millipore). To generate pPM09, pPM27, and pPM32, the *mEGFP*, *pHluorin*, and *mCherry* ORFs were amplified by PCR using pFa6a-mEGFP-kanMX6 (a gift from Julien Berro^74^, Addgene plasmid # 87023), pAS1NB (a gift from Mark Prescott^37^, Addgene plasmid # 71245), and pFa6a-3xHA-mCherry-natMX6, respectively. The amplicons were then assembled into pK3F (linearized with *Spe*I (New England Biolabs)) using the Gibson Assembly Master Mix (New England Biolabs). pPM28 was similarly generated by Gibson Assembly of the *pHluorin* amplicon into pPM32. To generate pPM31, the ORF for *hfYFP*^64^ was yeast codon optimized using the codon optimization tool from IDT and synthesized as a g-block flanked by *Spe*I restriction enzyme recognition sites (IDT). The g-block was solubilized in TE and Gibson Assembled into pK3F as described above. To generate pPM29, an amplicon containing the *HEH1* promoter, the coding sequence of a fragment of Heh1 encoding the first 479 amino acids, followed by the *ADH1* 3’UTR, was amplified by PCR using genomic DNA from PMCPL1082 as a template and primers with *Eag*I and *Xho*I restriction enzyme recognition sites. The amplicon was restriction digested with *Eag*I (New England Biolabs) and *Xho*I (New England Biolabs) and ligated (T4 DNA ligase, Invitrogen) into pRS405 digested with *Eag*I and *Xho*I. To generate pPM30, an ORF containing the mCherry coding sequence was generated by PCR using pSJ1321 (a gift from Sue Jaspersen^75^, Addgene plasmid #86413) as a template and primers with *BamH*I and *Pac*I restriction enzyme recognition sites. The amplicon was restriction digested with *BamH*I (New England Biolabs) and *Pac*I (New England Biolabs) and ligated into gel purified pPM29 digested with *BamH*I and *Pac*I.

pPM03 was generated in two steps. First, the genomic region containing the *ATG8* promoter (576 base pairs immediately upstream of the *ATG8* start codon) was generated by PCR using genomic DNA from W303A and primers with *Sac*I and *Eag*I restriction enzyme recognition sites. The amplicon was restriction digested with *Sac*I (New England Biolabs) and *Eag*I ligated into pRS406 linearized with *Sac*I and *Eag*I. Next, the genomic region containing the *ATG8* terminator (319 base pairs immediately downstream of the *ATG8* stop codon) was generated in a similar manner using primers with *Hind*III and *Sal*I restriction enzyme recognition sites. The amplicons were digested with *HindIII* and *SalI* ligated into the plasmid containing the *ATG8* promoter linearized with *Hind*III (New England Biolabs) and *Sal*I (New England Biolabs). To generate pPM11, the *DNM1* gene was amplified by PCR using genomic DNA from W303A and primers with *Eag*I and *Spe*I restriction enzyme recognition sites. The amplicon was restriction digested with *Eag*I and *Spe*I and ligated into pPM03 linearized with *Eag*I and *Spe*I.

To generate pPM15, site-directed mutagenesis of the codon encoding lysine at amino acid position 41 was altered to alanine using the Q5 site-directed mutagenesis kit (New England Biolabs) using pPM11 as a template. All plasmid sequences were confirmed by Sanger sequencing.

### RTqPCR

RNA extraction was performed using the MasterPure Yeast RNA purification kit (Lucigen) from samples treated as indicated in the legend of Figure S1. cDNA was produced from ∼1 µg of RNA using M-MuLV Reverse Transcriptase (New England Biolabs). RTqPCR was performed on 1:10 dilutions of the cDNA using iTaq Universal SYBR Green Supermix (Bio Rad) and primer pairs validated for RTqPCR efficiency for *ATG39* (sATG39 qPCR 6 and asATG39 qPCR 6), *ATG8* (sATG8 qPCR and asATG8 qPCR), *ATG11* (sATG11 qPCR and asATG11 qPCR), *ATG40* (sATG40 qPCR and asATG40 qPCR), and *ACT1* (sACT1 qPCR and asACT1qPCR) which served as a control. All PCRs were performed using CFX96 Touch Real-Time Detection System thermocycler (BioRad). Cq values as reported by the thermocycler during the RTqPCR for the gene of interest were subtracted from those of *ACT1* to obtain ΔCq. ΔΔCq was then obtained by subtracting the ΔCq value of the control condition from the condition of interest (i.e. rapamycin or nitrogen starvation). Fold change was calculated as 2^−*ΔΔCq*^.

### Live-cell Microscopy

For all live-cell imaging, cells incubated in sd-N for the duration of time indicated in the figure legend were gently collected by centrifugation and resuspended in sd-N medium prior to imaging. Images from Figure 1C, Figure S1F, Figure S2D, and the images used to generate the data in Figure 3D were acquired on a DeltaVision microscope (Applied Precision) equipped with a UPlanSapo x100 1.4 numerical aperture oil immersion objective (Olympus), and an AURA light engine (Lumencor) and CoolSnapHQ^2^ charge-coupled device (CCD; Photometrics) camera imaging using an Insight SSI four-color live-cell filter set (GFP: excitation, 425–495 nm and emission, 500–550 nm; YFP: excitation, 496–528 nm and emission, 537–559 nm; mCherry: excitation, 555–590 nm and emission, 600– 675 nm) using Resolve3D in SoftWoRx 7.0.0 software. The microscope stage and objectives were maintained at 30°C within an environmental chamber.

All other images were acquired on a lattice light sheet microscope^76^ equipped with a 25X 1.1 numerical aperture CFI APO LWD objective (Nikon) whose focal plane is coincident with the light sheet, and a sCMOS camera (Hamamatsu Orca Flash 4.0 v2). Cells were added to 5 mm round cover slips (Warner Instruments) coated with Concanavalin A (Sigma) as described in^77^. The coverslip with the cells adhered was then placed in an imaging chamber in a bath containing sd-N medium warmed to 30°C.

### Correlative light and electron microscopy

Correlative light and electron microscopy of resin-embedded cells was performed based on the protocol published by Kukulski and colleagues^78^. In brief, cells (PMCPL1283 and PMCPL1390) were cultured in sd-N for four hours. Cells were pelleted by centrifugation at ∼800 RCF for five minutes and high-pressure frozen in the 200-µm recess of an aluminum platelet (Engineering Office M. Wohlwend 241) using an HPM100 (Leica Microsystems). The samples were then freeze substituted in 0.1% uranyl acetate. Automated temperature control was then used to complete the solution exchange and embedding in Lowicryl HM20 (Polysciences). Sectioning was performed using a diamond knife (Diatome) mounted on an ultramicrotome (Leica Artos 3D) to a nominal thickness of 250 µm. Sections were then collected on carbon-supported 200-mesh copper grids (Ted Pella 01840).

The position of Atg39-hfYFP was determined by fluorescence microscopy. Next, 15 nm protein A-coated gold beads (Cell Microscopy Core, University Medical Center Utrecht) were adhered to the grids to aid in the tilt-series alignment.

We used automated tilt-series acquisition via SerialEM 3.6.15^79^ on an electron microscope (FEI TF20) operating at 200 kV from approximately −65 to +65 degrees (one-degree increments) with a high-tilt tomography holder (Fischione Instruments 2020), a 100 µm objective aperture, a 150 µm C2 aperture, and a 4k x 4k Eagle CCD (FEI) at a binned pixel size of 1.242 nm. Single- or dual-axis tilt series were acquired as necessary to achieve sufficient contrast to visualize membranes. IMOD 4.11.24^80^ was used for reconstruction (in an automated fashion^81^), nonlinear anisotropic diffusion filtering, and manual segmentation. Vph1-3xHA-mCherry and Nup170-mCherry fluorescence were used to correlate fluorescence and electron microscopy images to localize the Atg39-hYFP within the ultrastructure.

### Calculating expected time resolution in observable morphological intermediates in CLEM

A modification of the popular coupon collector’s problem^82^ from probability theory was used to estimate the number of tomograms needed to observe all morphologies in nucleophagy which persist for a given amount of time. The equation used is

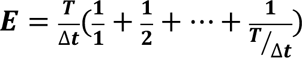

where E is the expected number of tomograms need to visualize at least one example of all morphological intermediates that exist for a time (Δt) as a ratio to the average time (T) Atg39-foci are outside of the vacuole (∼300 seconds, Figure 3I). Using this formula, we calculated that 30 tomograms were sufficient to achieve a time resolution of 30 seconds. Simply put, it is statistically likely to observe at least one instance of a morphological intermediate in nucleophagy that is identifiable by CLEM that persists for more than 30 seconds in a random selection of 30 tomograms.

### Image processing and analysis

Micrographs acquired on the Applied Precision DeltaVision microscope were deconvolved using an iterative algorithm in SoftWoRx (6.5.1; Applied Precision, GE Healthcare). Unprocessed images were used for all quantification of fluorescence intensity on micrographs acquired on the DeltaVision. Fluorescence intensity values from micrographs were measured in FIJI/imageJ^83^. Micrographs acquired on the LLSM were deskewed and deconvolved as described in^76^. LLSM images were rendered in 3D (Figure 3A and 3B) using the commercial software Imaris (Oxford Instruments).

### Nucleophagic flux calculations

Cells expressing Atg39-pHluorin-mCherry (PMCPL1045, PMCPL1078, PMCPL1079, PMCPL1253, PMCPL1130, PMCPL1282, PMCPL1066, PMCPL1249, PMCPL1276, PMCPL1243, PMCPL1252, PMCPL1264, PMCPL1067, PMCPL1127, PMCPL1293, PMCPL1274, PMCPL1124, PMCPL1421, PMCPL1430, PMCPL1443, PMCPL1493, PMCPL1496) were cultured overnight, incubated in sd-N media for 6 hours, and prepared for imaging on the DeltaVision. To calculate nucleophagic flux, all the Atg39-pHluorin-mCherry foci in the cell were manually counted and the number of red-only foci was divided by the sum of the red-only and red/green foci. The ratio obtained in WT was normalized to 1 and used to compare to all other genotypes.

### NE mCherry fluorescence calculation

Cells expressing Atg39-pHluorin-mCherry (PMCPL1045, PMCPL1078, PMCPL1079, PMCPL1253, PMCPL1130, PMCPL1282, PMCPL1066, PMCPL1249, PMCPL1276, PMCPL1243, PMCPL1252, PMCPL1264, PMCPL1067, PMCPL1127, PMCPL1293, PMCPL1274, PMCPL1124, PMCPL1421, PMCPL1430, PMCPL1443, PMCPL1493, PMCPL1496) were cultured overnight, incubated in sd-N media for 6 hours, and prepared for imaging on the DeltaVision. To measure NE mCherry fluorescence, a 4 pixel-wide ROI was drawn along the nuclear periphery in regions devoid of Atg39 foci and the mean fluorescence intensity was measured and background fluorescence was subtracted.

### Nucleophagy frequency quantification

The number of Atg39-GFP accumulations in an experiment (three biological replicates total) was divided by the total number of cells. The resulting number was then divided by the duration of the experiment to obtain the number of nucleophagic events/cell/hour.

### Determining Atg39-GFP copy number

Atg39-GFP expressing cells (PMCPL940, PMCPL856) were incubated in sd-N media for four hours prior to imaging and each were separately mixed with Fta3-GFP expressing cells (MKSP3186) and mounted onto coverslips. Raw images were then analyzed using a custom-written MATLAB function, ‘findIrregularSpots3’ described previously^60^ that finds fluorescent objects regardless of their shape and returns xyz-coordinates of the object centroids. The xyz coordinates were then used as inputs for a custom-written MATLAB function ‘fitSpots3’ to fit and compute the volume of a 2D Gaussian to a Z-slice image closest to the z-coordinate of the spot centroid. Fta3-GFP spots were manually curated for their position in the cell cycle and those that were determined to be in anaphase B were selected for analysis. Atg39-GFP copy number was estimated by dividing the Gaussian volume for each spot by the average Gaussian volume of the Fta3-GFP spots, previously calculated to be 111 molecules^63^. Fta3-GFP foci identified as outliers using ROUT test Q=1%, one iteration, were excluded.

### Particle tracking/colocalization

Cells (PMCPL940, PMCPL857, PMCPL831, PMCPL832, PMCPL840, PMCPL1319, PMCPL1320, and PMCPL1087) were incubated in sd-N media for three hours before being prepared for imaging on the LLSM. Cells where a Atg39-GFP focus appeared that underwent nucleophagy were selected for further processing using MATLAB scripts to detect Atg39-GFP foci (spots) as described in “Determining Atg39-GFP copy number.” For particle tracking and colocalization analysis the spots were organized into trajectories using previously described MATLAB implementation^61^ of the widely-used tracking algorithm ‘track.m’^62^. Trajectories of a single Atg39-GFP spot that were erroneously split in two (when displacement between two frames exceeded a maximum linking distance) were linked together manually. For the relative spot intensity analysis, Gaussian volumes were calculated from Atg39-GFP spots as described above. The Gaussian volume of the tracked spot was normalized to that of the first time point in the trajectory. The spot coordinates in each track were also used to calculate displacement of Atg39-GFP between frames using a custom Jupyter Notebook script, implemented in Python (Figure 3E,G, Figure S2C).

For colocalization analysis, a custom Jupyter Notebook Python script used the coordinates of the identified Dnm1-mCherry spots which correspond to the time points of the Atg39-GFP tracks to calculate the distance from the center of the Atg39-GFP spots to the center of the closest Dnm1-mCherry spot (Figure 2D,E, Figure S2A,B). For all time points in which the distance between Atg39-GFP and Dnm1-mCherry spots were calculated to be 400 nm or less, a small ROI around the Atg39-GFP spot was cropped and the Pearson’s correlation coefficient between the two channels was determined using the Fiji plugin JaCoP^84^. The particles were considered colocalized if the Pearson’s correlation coefficient was >0.5.

All MATLAB and Jupyter Notebook Python script are available on GitHub.

### Quantification of INMDV diameter

The diameter of the INMDVs was measured in IMOD.

### Estimation of the percentage of INM turned over by nucleophagy in 24 hours

To estimate the percentage of INM that could be turned by nucleophagy in 24 hours, the amount of surface area of the INMDV removed in one round of nucleophagy in WT cells was determined. For this, the surface area of INMDVs in category 2-4 (i.e. after release as cytosolic NDVs) were approximated as ellipsoids and the length of the major and minor axes were measured on IMOD. The a and b axes (major and minor axes) were measured as the maximum distance within the membrane of the INMDV and the distance within the membrane orthogonal to the maximum, respectively. The c axis was recorded as the distance between the intersection of the major and minor axes and the edge of the INMDV. The surface area was then approximated using the a, b, and c radii and the ellipsoid surface area formula. The median value was then multiplied by the frequency of nucleophagy (1.4 events/cell/hour) and by 24 hours to estimate the surface area of the of INM removed per cell in 24 hours. This value was calculated as ∼7.14 µm^2^. To estimate the surface area of the NE prior to nitrogen starvation, cells expressing Heh1-GFP were imaged in log phase, and the radii of the nuclei were measured, and the surface area was calculated using the equation for the surface area of a sphere. The mean value was determined to be ∼8.63 µm^2^.

### Statistical methods

Graphs were generated using Prism (GraphPad 9.0). Where relevant, we used an appropriate experimental group that was carrier treated as a reference to test for contribution of other covariates. Statistical significance tests were used as indicated in figure legends. Significance values were calculated within Prism (GraphPad 10.0) and p-values are indicated on the graph or in figure legends as: *ns*, *p* > 0.05; * *p* ≤ 0.05; ** *p* ≤ 0.01; *** *p* ≤ 0.001; **** *p* ≤ 0.0001. All data was assumed to be normal with equal variance. Error bars are described in figure legends.

### Whole cell extract preparation

5 O.D. cells were pelleted by centrifugation and resuspended in 10% TCA for 1 h on ice and centrifuged at 15,000 *g* for 10 min at 4°C. The pellet was washed with ice-cold acetone, homogenized by sonication (Bioruptor UCD-200) and pelleted by centrifugation. After two cycles of washing and sonication, the pellet was vacuum-dried for 15 min. The dried cell pellet was then mechanically disrupted with 100 µl glass beads (Sigma) and 100 µl 50 mM Tris-HCl pH 7.5, 8 M urea, 2% SDS, and 1 mM EDTA, followed by addition of 100 µl protein sample buffer (Tris-HCl pH 6.8, 7 M urea, 10% SDS, 24% glycerol, bromophenol blue, and 10% β-mercaptoethanol).

### Immunoblotting

Proteins from whole cell extracts were resolved on 4-20% SDS-polyacrylamide gels (BioRad), followed by transfer of the proteins to 0.2 µm nitrocellulose membranes (Bio-Rad). The membranes were blocked in 5% non-fat milk in TBST for 1 h and immunoblotted with an antibody against HA directly conjugated to HRP (anti-HA-HRP; Roche) for 1 h at room temperature or with an antibody against GFP (anti-GFP; Biolegend) overnight at 4°C. Blots incubated with anti-GFP were then incubated with a secondary antibody against mouse (anti-mouse-HRP; Invitrogen) for 1 h at room temperature. Blots were visualized by ECL (Thermo Fisher Scientific) using a VersaDoc Imaging System (BioRad). Relative protein loading was visualized using Ponceau S Solution (Sigma).

## Data Availability

Source data will be provided with this paper at the time of publication. All other data supporting the findings presented in this study are available from the corresponding author upon reasonable request.

## Code Availability

The MATLAB and Jupyter Notebook Python scripts are available with no access restrictions at https://github.com/LusKingLab/.

## Supplemental Figures

**Supplemental Figure 1.**
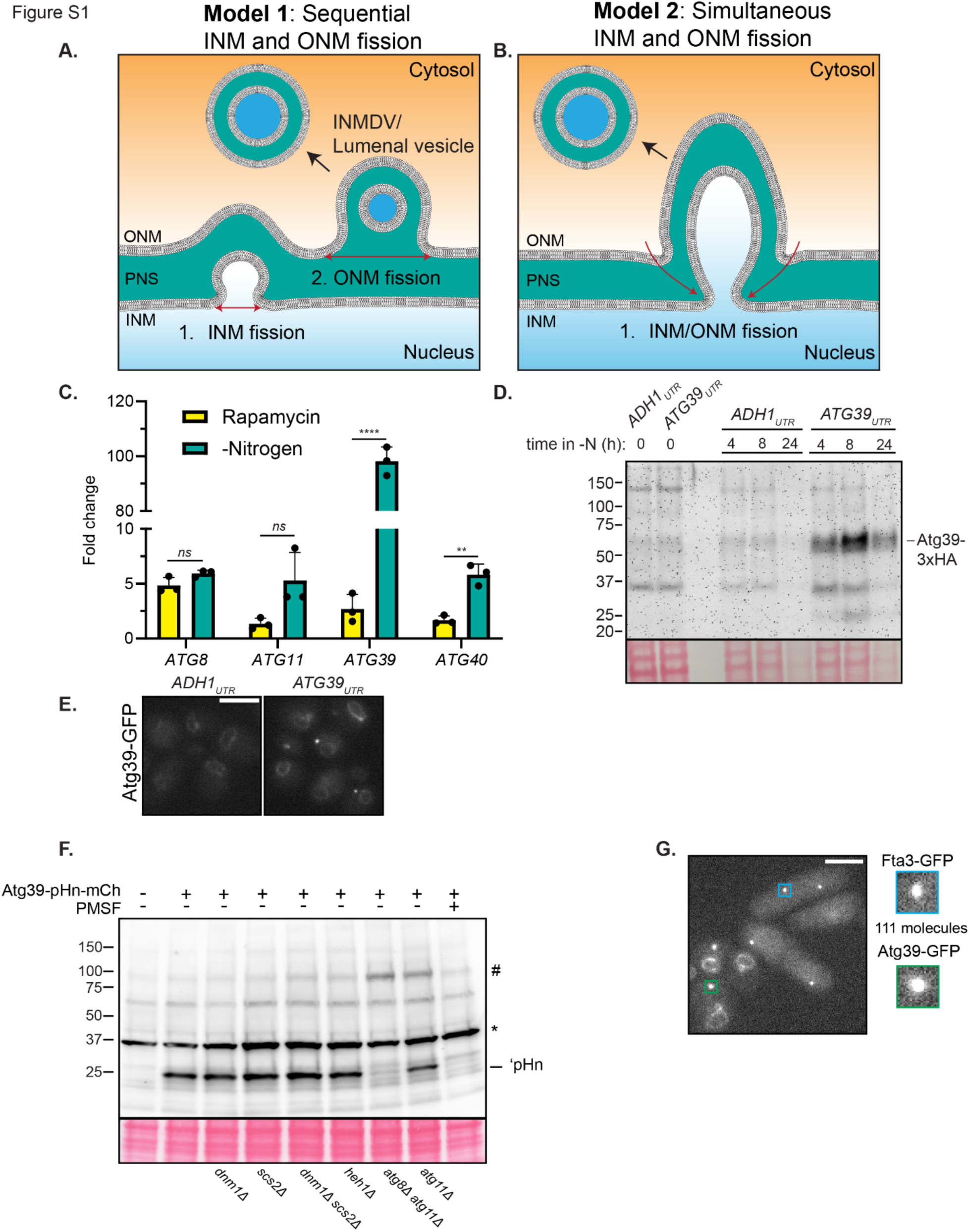
– Models of nucleophagy and characterization of Atg39 expression and copy number. **A,B)** Schematic of two models of nucleophagy incorporating either sequential (Model 1) or simultaneous (Model 2) INM and ONM fission events. Model 1 would also lead to the formation of a lumenal vesicle/INMDV intermediate. ONM, outer nuclear membrane; INM, inner nuclear membrane; PNS, perinuclear space; INMDV, inner nuclear membrane-derived vesicle; NDV, nucleus-derived vesicle. **C)** RTqPCR on indicated autophagy gene transcripts was performed in WT cells either treated with rapamycin for 2 h or cultured in medium lacking nitrogen for 2 h. Results displayed as a bar graph comparing the mean fold change of the transcripts between the rapamycin and -Nitrogen condition. Mean and standard deviation (error bars) of three biological replicates (black dots). Unpaired two-sided t tests. ****P<0.0001; **P<0.01; *ns*, not significant. **D)** Protein levels of Atg39-3xHA fusions derived from transcripts including either the *ADH1* 3’UTR or the endogenous *ATG39* 3’UTR were assessed by western blot with anti-HA antibody directly coupled to HRP (α-HA-HRP) and ECL. Positions of molecular weight standards (kD) at left. At bottom, membranes were stained with Ponceau to evaluate total protein loads. **E)** Fluorescence micrographs of Atg39-GFP produced from transcripts with 3’UTRs from the *ADH1* or *ATG39* genes after 4 h in medium lacking nitrogen. Scale bar is 5 µm. **F)** Western blot of whole protein extracts from cells expressing Atg39-pHn-mCh with fall out fragment (‘pHn) in the indicated genetic backgrounds and PMSF treatment after 24 h in sd-N. Position ‘pHn indicated on the right. An anti-GFP antibody and anti-mouse-HRP secondary followed by ECL was used to detect proteins. Positions of molecular weight standards (kD) at left. * is a non-specific band. # is the expected position of Atg39-pHn-mCh, which is undetectable in most backgrounds due to its low abundance and lack of sensitivity of the anti GFP antibody. **G)** To calculate copy number of Atg39-GFP within individual foci, we imaged budding yeast expressing Atg39-GFP next to *S. pombe* expressing Fta3-GFP, a kinetochore protein which has 111 molecules/focus in anaphase B^60^. By directly relating the fluorescence intensities, we calculated Atg39-GFP copy number/focus as presented in Figure 3D.

**Supplemental Figure 2.**
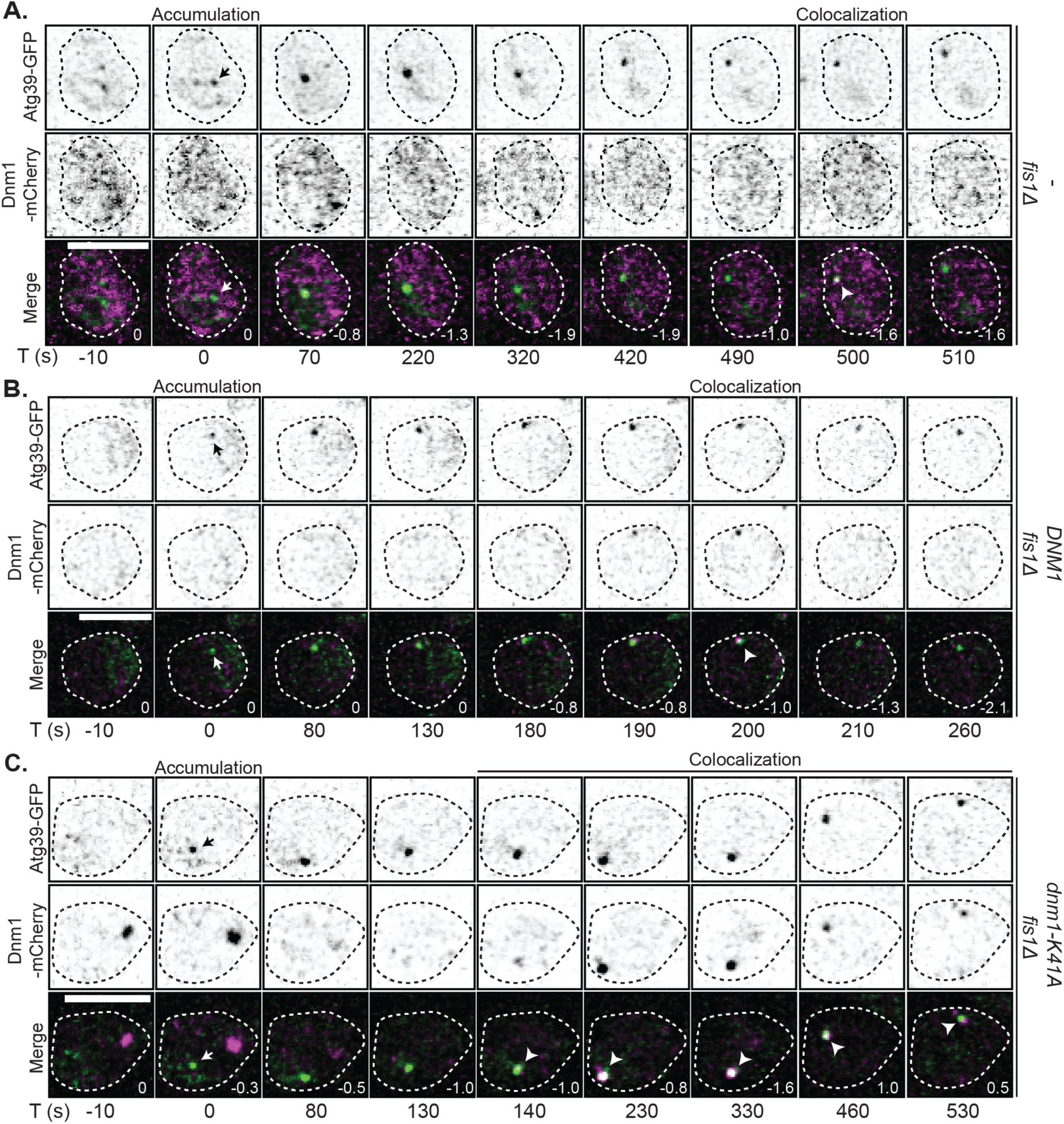
– Dnm1 colocalizes with Atg39 in the absence of Fis1. **A-C)** Inverted fluorescence micrographs of a timelapse series of *fis1Δ* cells expressing Atg39-GFP and Dnm1-mCherry with and without either an extra copy of *DNM1* or the dominant negative *dnm1-K41A* allele under conditions of nitrogen starvation. Green and red channels shown with merge at indicated times (T) with T=0 s being the first detection of a Atg39-GFP focus (arrow). Arrowheads point to colocalization between Atg39-GFP and Dnm1-mCherry. Numbers in merge are the z-position (in μm) of the image shown in reference to the midplane. Scale bars are 5 µm.

**Supplemental Figure 3.**
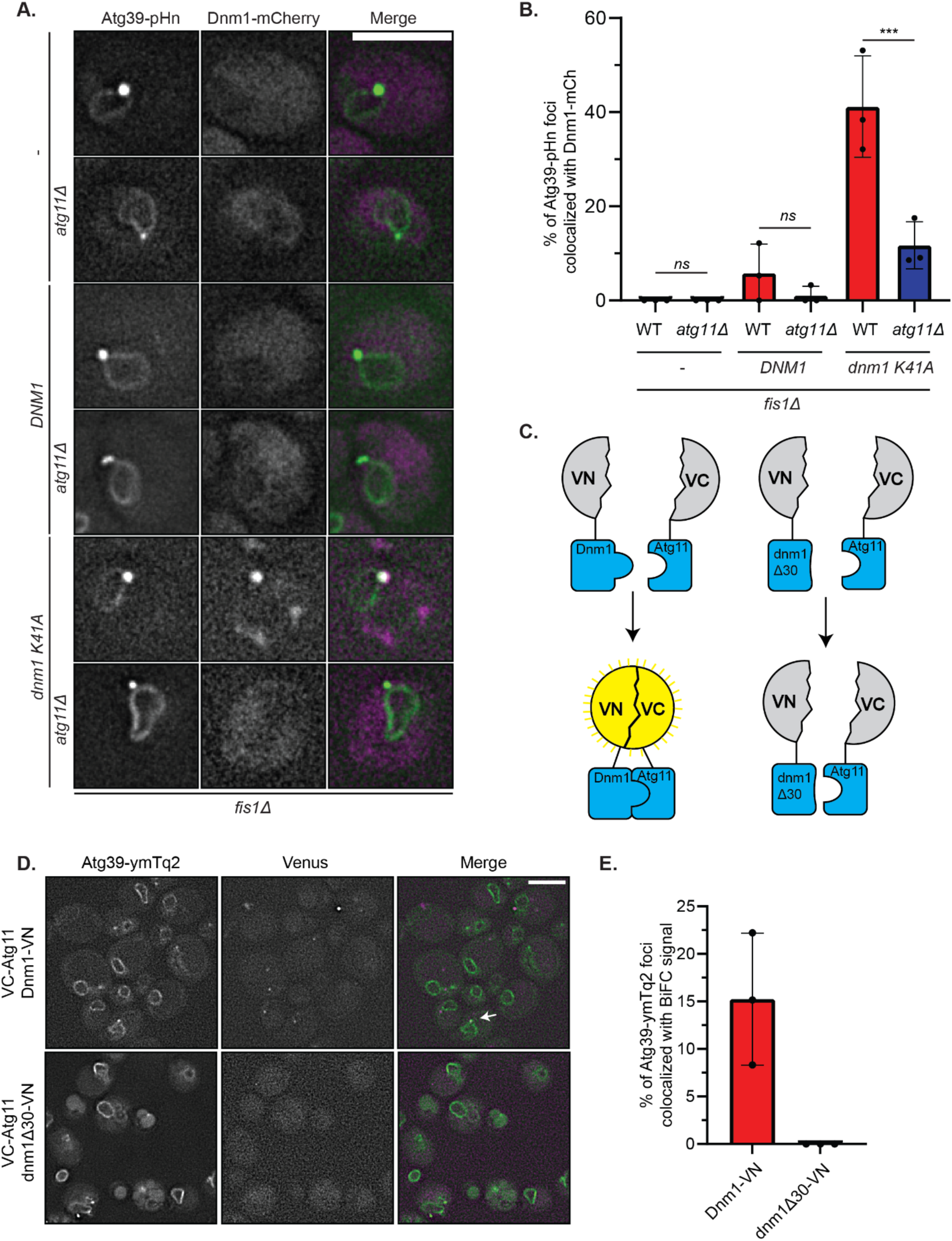
– Atg11 recruits Dnm1 to the NE. **A)** Fluorescence micrographs of *fis1Δ* cells and *fis1Δatg11Δ* cells expressing Atg39-pHn, Dnm1-mCherry with and without an extra copy of *DNM1* or the *dnm1-K41A allele* under conditions of nitrogen starvation. Green and red channels shown with merge. Scale bar is 5 µm. **B)** Plot of the mean percentage of Atg39-pHn foci that colocalize with Dnm1-mCherry from experiments in **A**. Mean and standard deviation (error bars) from three biological replicates (black dots). For *fis1Δ* cells, *n*=79 Atg39-pHn foci. For *fis1Δ* cells expressing *DNM1*, *n*=104 Atg39-pHn foci. For *fis1Δ* cells expressing *dnm-K41A*, *n*=166 Atg39-pHn foci. For *fis1Δ atg11Δ* cells, *n*=78 Atg39-pHn foci. For *fis1Δatg11Δ* cells expressing *DNM1*, *n*=96 Atg39-pHn foci. For *fis1Δatg11Δ* cells expressing *dnm1-K41A*, *n*=165 Atg39-pHn foci. Unpaired two-sided t test. ***P<0.001; *ns*, not significant. **C)** Schematic of BiFC. VN and VC are the N- and C-terminal fragments of Venus, respectively. An interaction between Dnm1-VN (but not dnm1Δ30) and Atg39-VC leads to the reconstitution of Venus and its fluorescence. **D)** Fluorescence micrographs of cells expressing Atg39-ymTurquoise2 (Atg39-ymTq2), VC-Atg11, and either Dnm1-VN or dnm1Δ30-VN under conditions of nitrogen starvation. Yellow (magenta) and blue (green) channels shown with merge. The arrow points out colocalization between the Venus and Atg39-ymTq2 foci. Scale bar is 5 µm. **E)** Plot of the mean percentage of Atg39-ymTq2 foci that colocalize with Venus foci from experiments in **D**. Mean and standard deviation (error bars) from three biological replicates. For cells expressing Dnm1-VN, *n*=84 Atg39-ymTq2 foci. For cells expressing dnm1Δ30-VN, *n*=93 Atg39-ymTq2 foci.

**Supplemental Figure 4.**
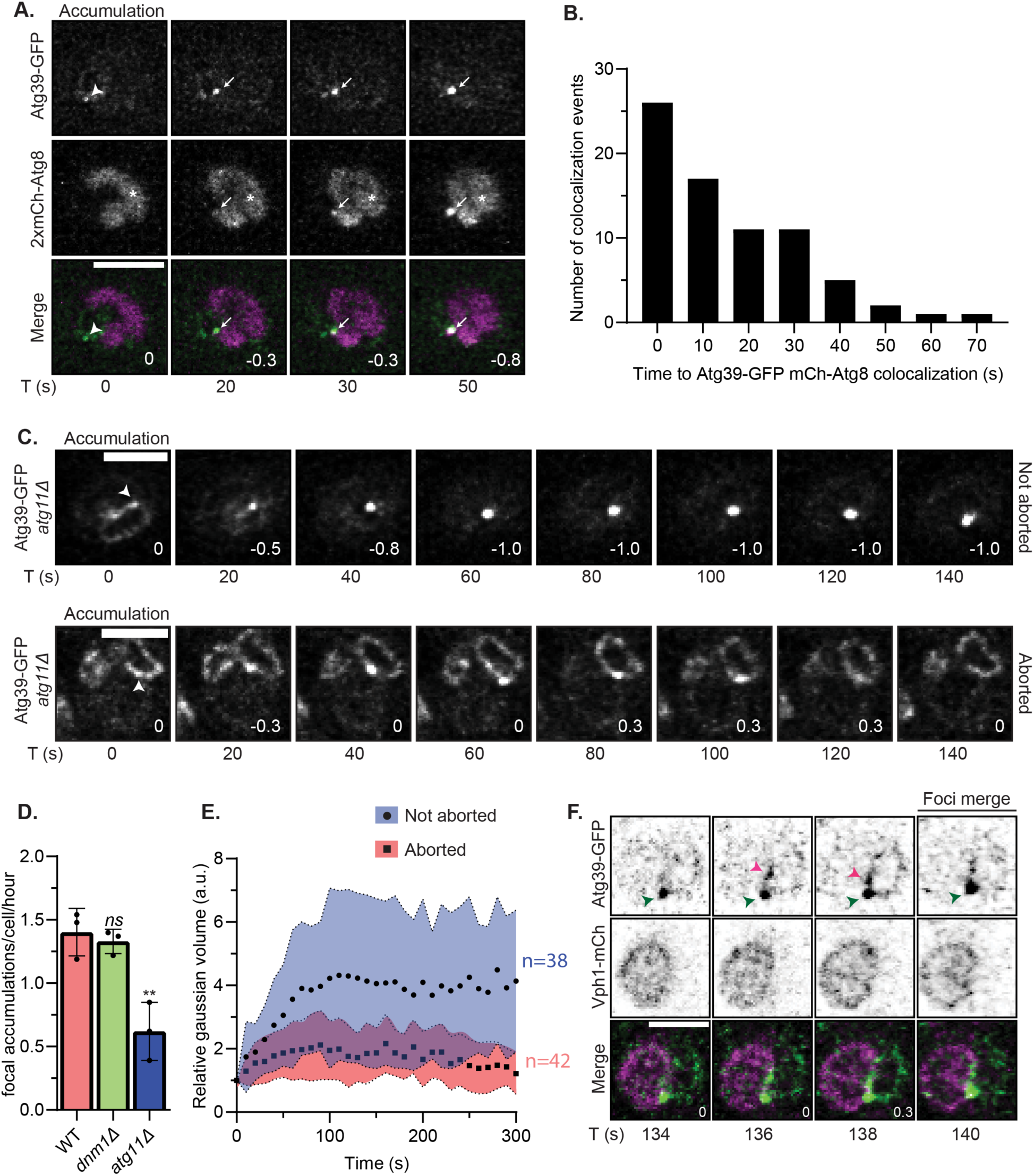
– Atg8 recruitment, Atg11 dependence, and Atg39 dynamics during nucleophagy. **A)** Fluorescence micrographs of cells expressing Atg39-GFP and 2xmCherry(mCh)-Atg8 imaged every 10 seconds under conditions of nitrogen starvation. Green and red channels shown with merge. Numbers in merge are z-position (in μm) of the displayed image in reference to the midplane. T (s) indicates the time in seconds since the first detection of the Atg39-GFP focus (indicated above, arrowhead). Arrows point to colocalization between Atg39-GFP and 2xmCh-Atg8. * indicates fluorescence from the vacuole in the red channel. Scale bar is 3 µm. **B)** Histogram of the number of colocalization instances for the indicated time after the appearance of an Atg39-GFP focus. Data are from the experiments in **A**; *n*=73 cells, 74 tracks, three biological replicates. **C)** Fluorescence micrographs of *atg11Δ* cells expressing Atg39-GFP, Vph1-mCherry, and Nup170-mCherry every 10 seconds under conditions of nitrogen starvation. Green channel is shown. Numbers in merge are z-position (in μm) of the displayed image in reference to the midplane. T (s) indicates the time in seconds since the first detection of the Atg39-GFP focus (indicated above, arrowhead). Cells shown are representative examples from the two groups of tracks in which the Atg39-GFP focus disappears prior to internalization in the vacuole (aborted) or not (not aborted). **D)** Plot of the mean number of focal accumulations divided by the total number of cells divided by the total time of the experiment (h) for the indicated genotype. Data are from three biological replicates (black dots). For WT cells, *n*=294 cells. For *dnm1Δ* cells, *n*=557 cells. For *atg11Δ* cells, *n*=522 cells. Ordinary one-way ANOVA with multiple comparisons with **P<0.01; *ns*, not significant. **E)** Fluorescence intensity of Atg39-GFP foci in *atg11Δ* cells plotted versus time after their first detection. Data are split into two groups, aborted (red, squares) and not aborted (blue, circles) from three biological replicates. For cells in the aborted group, *n*=40 cells, 42 tracks. For cells in the not aborted group, *n*=37 cells, 38 tracks. **F)** Inverted fluorescence micrographs of WT cells expressing Atg39-GFP and Vph1-mCherry (mCh) imaged every two seconds under conditions of nitrogen starvation. Green and red channels shown with merge. Numbers in merge are z-position (in μm) of the displayed image in reference to the midplane. T (s) indicates the time in seconds since the first detection of the Atg39-GFP focus (green arrowhead). Another less intense focus of Atg39-GFP is visible merging with the initial particle (pink arrowhead). Scale bar is 3 µm.

**Supplemental Figure 5.**
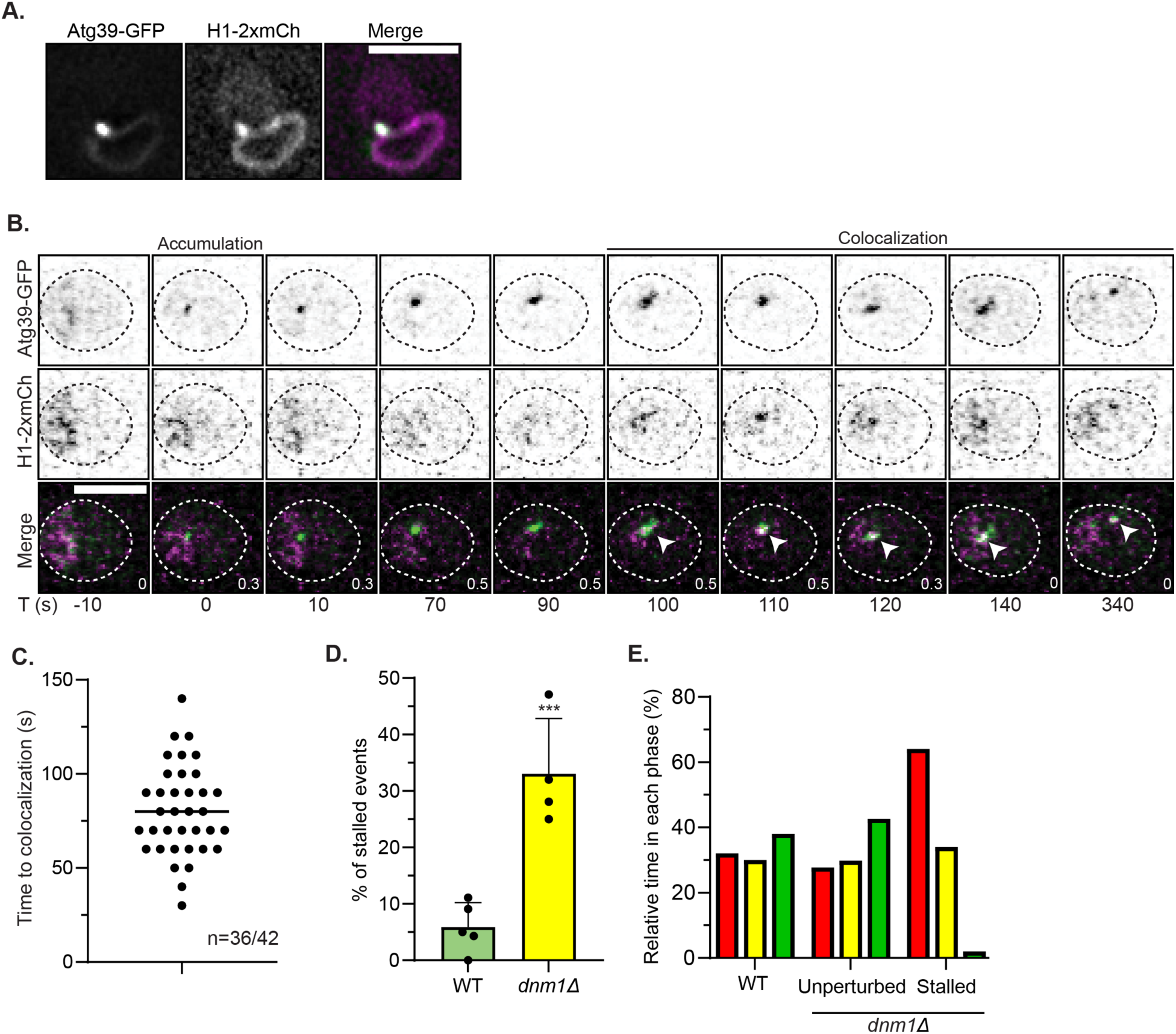
– Characteristics of Atg39-GFP mobility and cargo capture. **A)** Fluorescence micrographs of WT cells expressing Atg39-GFP and a model INM cargo comprising the INM targeting and transmembrane domain of Heh1 fused to two tandem mCherry fluorophores (H1-2xmCh) under conditions of nitrogen starvation. Green and red channels shown with merge. Scale bar is 3 µm. **B)** Inverted fluorescence micrographs of WT cells expressing Atg39-GFP and H1-2xmCh under conditions of nitrogen starvation. Imaged every 10 seconds with time 0 assigned to the point of Atg39-GFP focal accumulation. Green and red channels shown with merge. Arrowheads point to colocalization between Atg39-GFP and H1-2xmCh. Cell boundaries are outlined. Numbers in merge are z-position (in μm) of the displayed image in reference to the midplane. Scale bar is 3 µm. **C)** Scatter plot depicting the elapsed time from the first detection of an Atg39-GFP focus and the first instance of colocalization with H1-2xmCh from experiments as in **B**. The median is depicted by a black bar. *Data are from n=36 cells in which Atg39-GFP and H1-2xmCh were found colocalized outside of the nucleus out of n=42 total cells from two biological replicates*. **D)** Bar graph of the percentage of Atg39 foci tracked from first appearance that never make it into the vacuole over a 20-minute timelapse experiment as noted in Figure 3F for the indicated genotype. Mean and standard deviation (error bars) from 5 (WT cells) or 4 (*dnm1Δ* cells) biological replicates (black dots). Unpaired two-sided t test, *n*=114 tracks, 99 cells for WT cells and *n*=95 tracks, 82 cells for *dnm1Δ* cells. ***P<0.001. **E)** Plot of the fraction of time (as a percentage) that Atg39-GFP particles spend in each phase as per color code in Figure 3E and 3G in WT and *dnm1Δ* cells. The latter are also segregated based on those that enter the vacuole (unperturbed) and those that don’t (stalled).

**Supplemental Figure 6.**
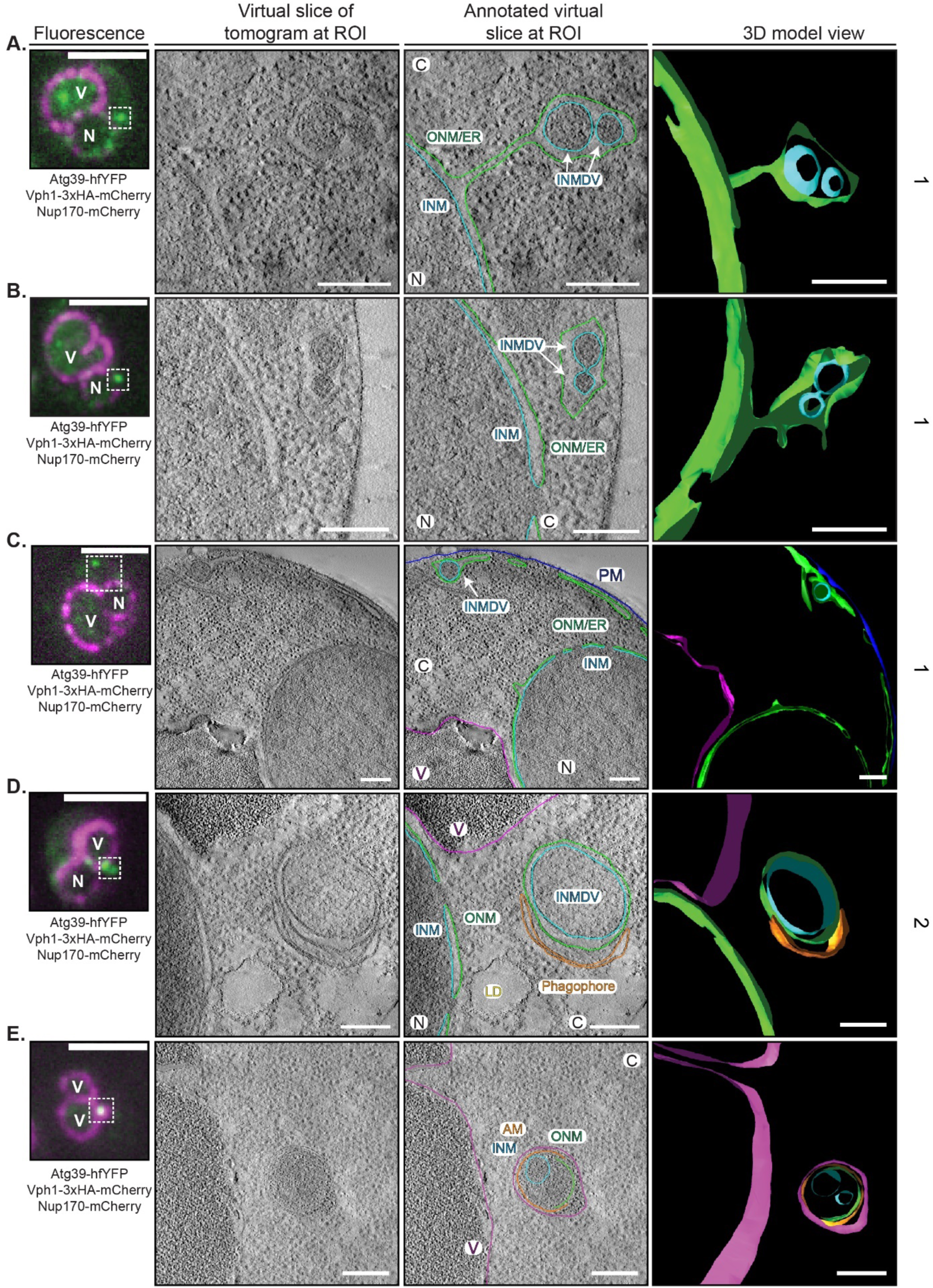
– Ultrastructure of sites of Atg39 accumulation during nucleophagy. **A-E)** CLEM/tomography of sites of Atg39-hfYFP foci. From left to right: fluorescence image of WT cells expressing Atg39-hfYFP (green), Vph1-mCherry and Nup170-mCherry (magenta). Scale bar is 3 µm. EM tomogram acquired of boxed region in fluorescence image and tomogram virtual slice without and with annotation shown alongside a snapshot of a 3D model. Arrows point to INMDVs. ONM is green, INM is light blue, phagophore/autophagosome is orange, vacuole membrane is magenta. Scale bar is 200 nm. C, cytosol; N, nucleus; V, vacuole; ONM, outer nuclear membrane; INM, inner nuclear membrane; ER, endoplasmic reticulum; INMDV, inner nuclear membrane derived vesicle; AM, autophagic membrane. Numbers at right refer to classification scheme in 4F.

**Supplemental Figure 7.**
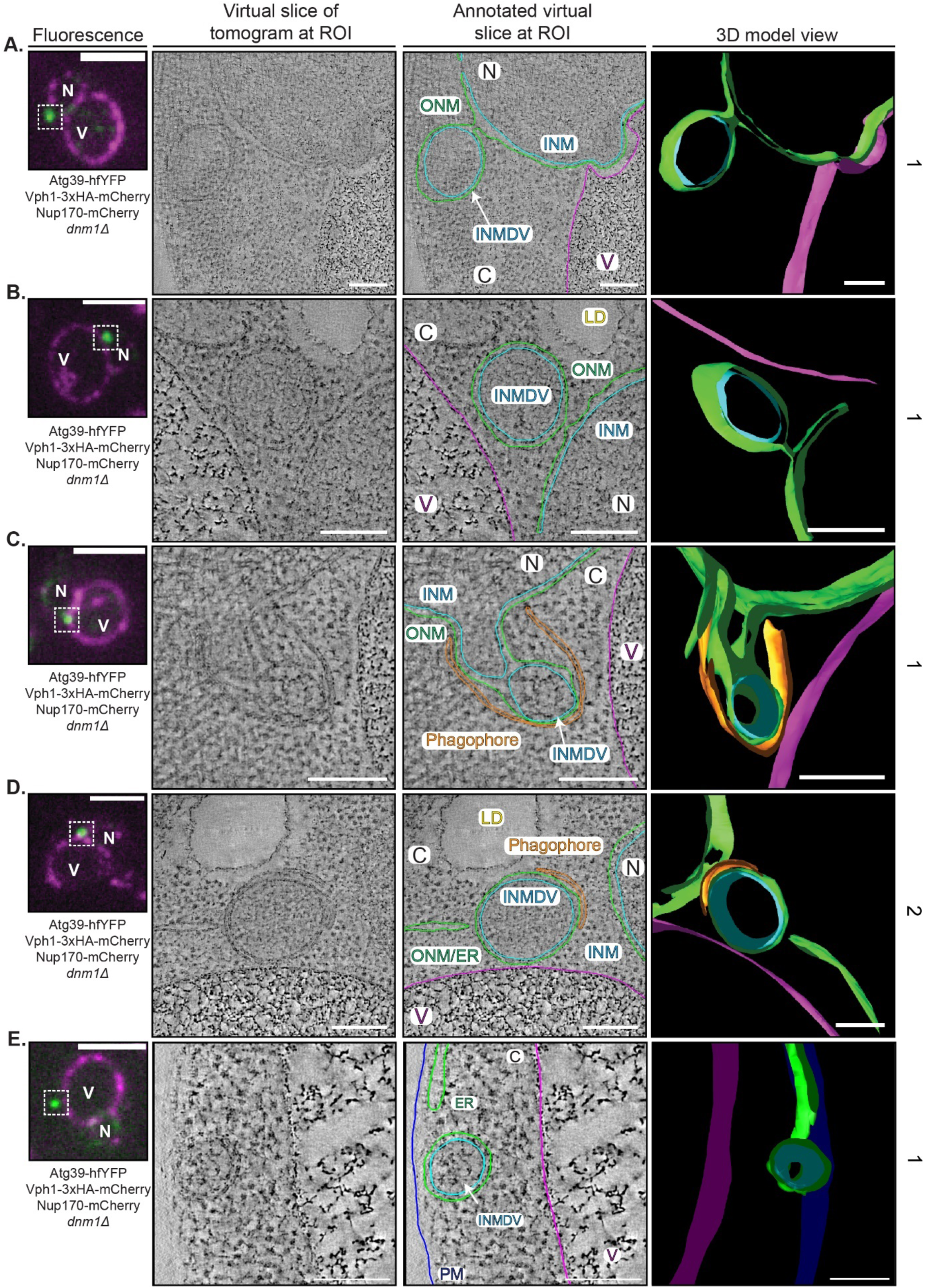
– Ultrastructure of sites of Atg39 accumulation in *dnm1Δ* cells. **A-E)** CLEM/tomography of sites of Atg39-hfYFP foci in *dnm1Δ* cells. From left to right: fluorescence image of WT cells expressing Atg39-hfYFP (green), Vph1-mCherry and Nup170-mCherry (magenta). Scale bar is 3 µm. EM tomogram acquired of boxed region in fluorescence image and tomogram virtual slice without and with annotation shown alongside a snapshot of a 3D model. Arrows point to INMDVs. ONM is green, INM is light blue, phagophore/autophagosome is orange, vacuole membrane is magenta. Scale bar is 200 nm. C, cytosol; N, nucleus; V, vacuole; ONM, outer nuclear membrane; INM, inner nuclear membrane; ER, endoplasmic reticulum; INMDV, inner nuclear membrane derived vesicle; AM, autophagic membrane; LD, lipid droplet. Numbers at right refer to classification scheme in Figure 4F.

**Supplemental Figure 8.**
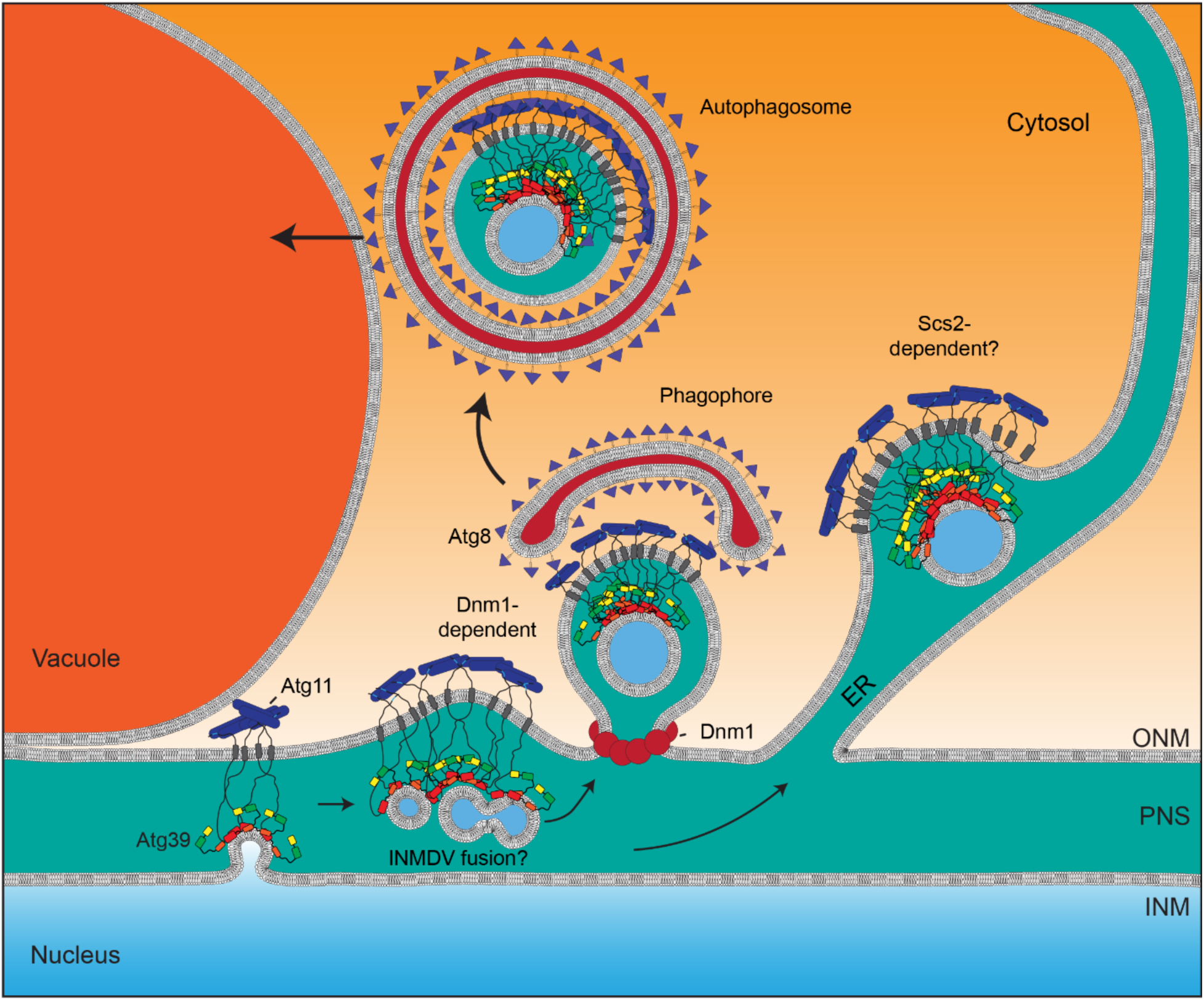
– Proposed model of nucleophagy. **A)** Illustrative model of the molecular and morphological steps of nucleophagy. Atg39 locally accumulates adjacent to the NVJ in a mechanism supported by Atg11 (blue cylinders). It is likely that this clustering coincides with cargo capture and the evagination and fission of the INM to generate an INMDV. Initial small (50-100nm INMDVs) may grow by fusion to generate larger (200-300 nm) INMDVs that are ultimately released from the NE through a Dnm1-dependent ONM fission step, or from the cortical ER through a proposed Scs2-dependent mechanism. Engagement with the phagophore occurs at or just after ONM fission. Autophagosomes containing double membrane nuclear derived vesicles (NDVs) fuse with vacuoles. Some INMDVs may also be capable of transiting through the broader ER but their fate remains uncertain.

## Movies

**Movie 1. LLSM movie of Atg39-GFP dynamics in WT cells.** A 5-minute timeseries of cells expressing Atg39-GFP and Vph1-mCherry imaged every 10 seconds by LLSM. Vph1-mCherry signal is rendered as a surface (magenta) and Atg39-GFP foci are rendered as balls (green). Scale bar is 10 µm. Frame rate is 24 frames per second.

**Movie 2. CLEM and tomography of potential interface between NDV in an autophagosome and the vacuole in a WT cell.** Electron tomogram of a cell expressing Atg39-hfYFP, Nup170-mCherry, and Vph1-mCherry obtained at a site of Atg39-hfYFP accumulation shown in Figure 4D. Note that the outer membrane of the autophagosome and the vacuole extend towards each other but do not touch. However, content with similar electron density of the vacuole lumen is observed between the inner and outer membrane of the autophagosome suggesting prior fusion with the vacuole. NE, green; INMDV, blue; autophagic membranes, orange; vacuole, magenta. Scale bar is 100 nm.

**Movie 3. CLEM and tomography of an INMDV within the NE/ER lumen engaged with a phagophore in a *dnm1Δ* cell.** Electron tomogram of a cell expressing Atg39-hfYFP, Nup170-mCherry, and Vph1-mCherry in a *dnm1Δ* cell obtained at a region of Atg39-hfYFP accumulation shown in Figure 5D. NE, green; INMDV, blue; autophagic membranes, orange; vacuole, magenta. Scale bar is 100 nm.

## Supplementary Tables

**Supplementary Table 1:**
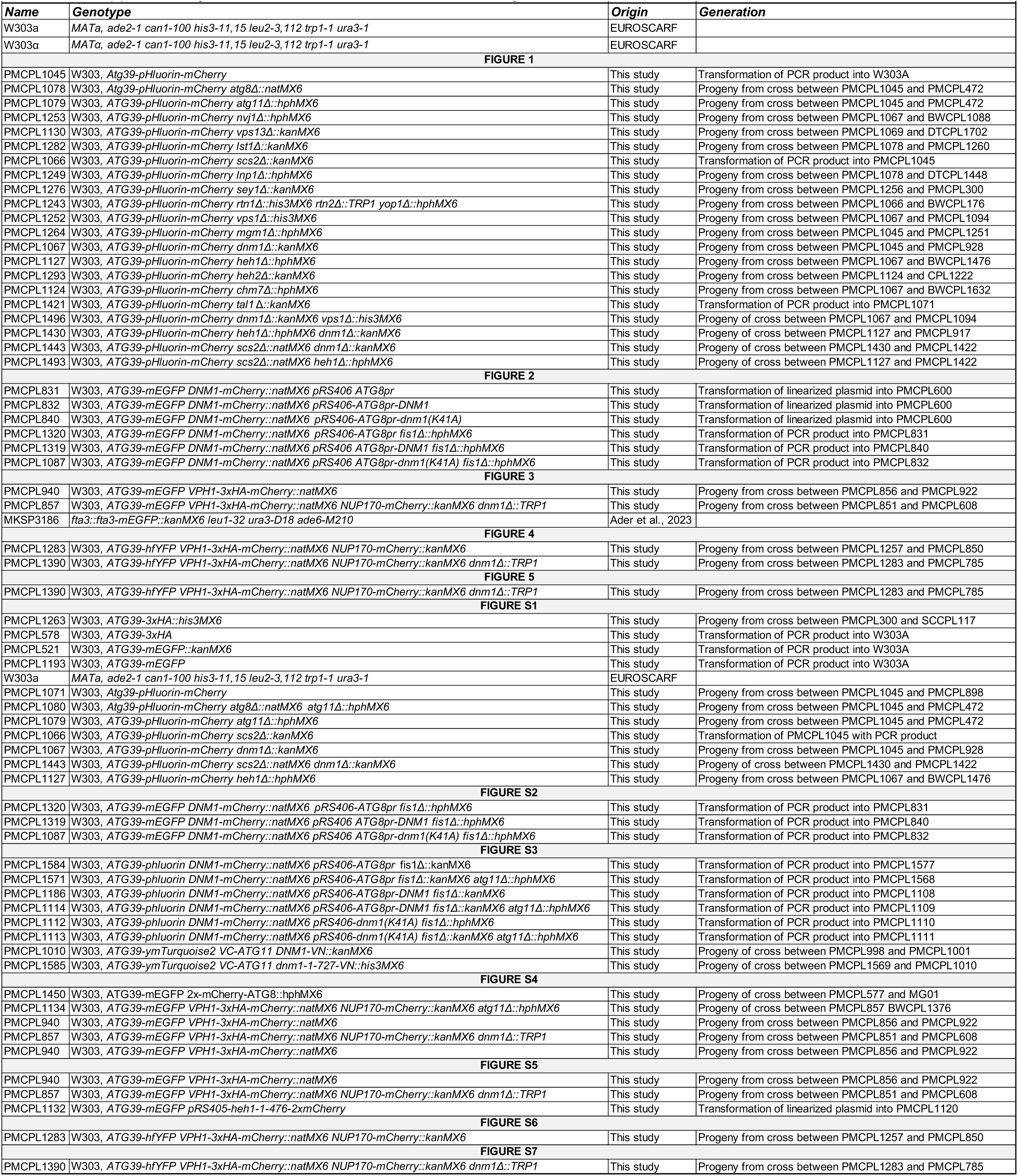
Yeast strains used in this study.

**Supplementary Table 2:** Plasmids used in this study.

| **Name** | **Description** | **Origin** |
| --- | --- | --- |
| pFA6a-mEGFP-kanMX6 | Template for PCR based chromosomal integration of mEGFP ORF | Lacy et al. 2017 |
| pFA6a-hphMX6 | Template for PCR based chromosomal integration of hphMX6 cassette | Longtine et al. 1998 |
| pFA6a-natMX6 | Template for PCR based chromosomal integration of natMX6 cassette | Van Driessche et al. 2005 |
| pFA6a-kanMX6 | Template for PCR based chromosomal integration of kanMX6 cassette | Bahler J. et al. 1998 |
| pAS1NB | Template for cloning pPM27 | Rosado et al. 2008 |
| pSJ1321 | Template for cloning pPM30 | Smoyer et al. 2016 |
| pFA6a-3xHA-mCherry-natMX6 | Template for PCR based chromosomal integration of 3xHA-mCherry ORF | This study |
| pFA6a-3xHA-his3MX6 | Template for PCR based chromosomal integration of 3xHA | Longtine et al. 1998 |
| pPM09 | mEGFP-loxP-kanMX6-loxP | This study |
| pPM26 | 3xHA-loxP-kanMX6-loxP | This study |
| pPM31 | hfYFP-loxP-kanMX6-loxP | This study |
| pPM27 | pHlourin-loxP-kanMX6-loxP | This study |
| pPM28 | pHlourin-mCherry-loxP-kanMX6-loxP | This study |
| pFA6a-mCherry-kanMX6 | Template for PCR based chromosomal integration of mCherry ORF | This study |
| pPM03 | pRS406 ATG8pr | This study |
| pPM11 | pRS406 ATG8pr-*DNM1* | This study |
| pPM15 | pRS406 *dnm1* (*K41A*) | This study |
| pSH47 | pRS416 *GAL1-CRE* | Güldener et al. 1996 |
| pPM29 | pRS405 heh1-1-479-mCherry | This study |
| pPM30 | pRS405 heh1-1-479-2xmCherry | This study |
| pPM32 | mCherry-loxP-kanMX6-loxP | This study |

**Supplementary Table 3:**
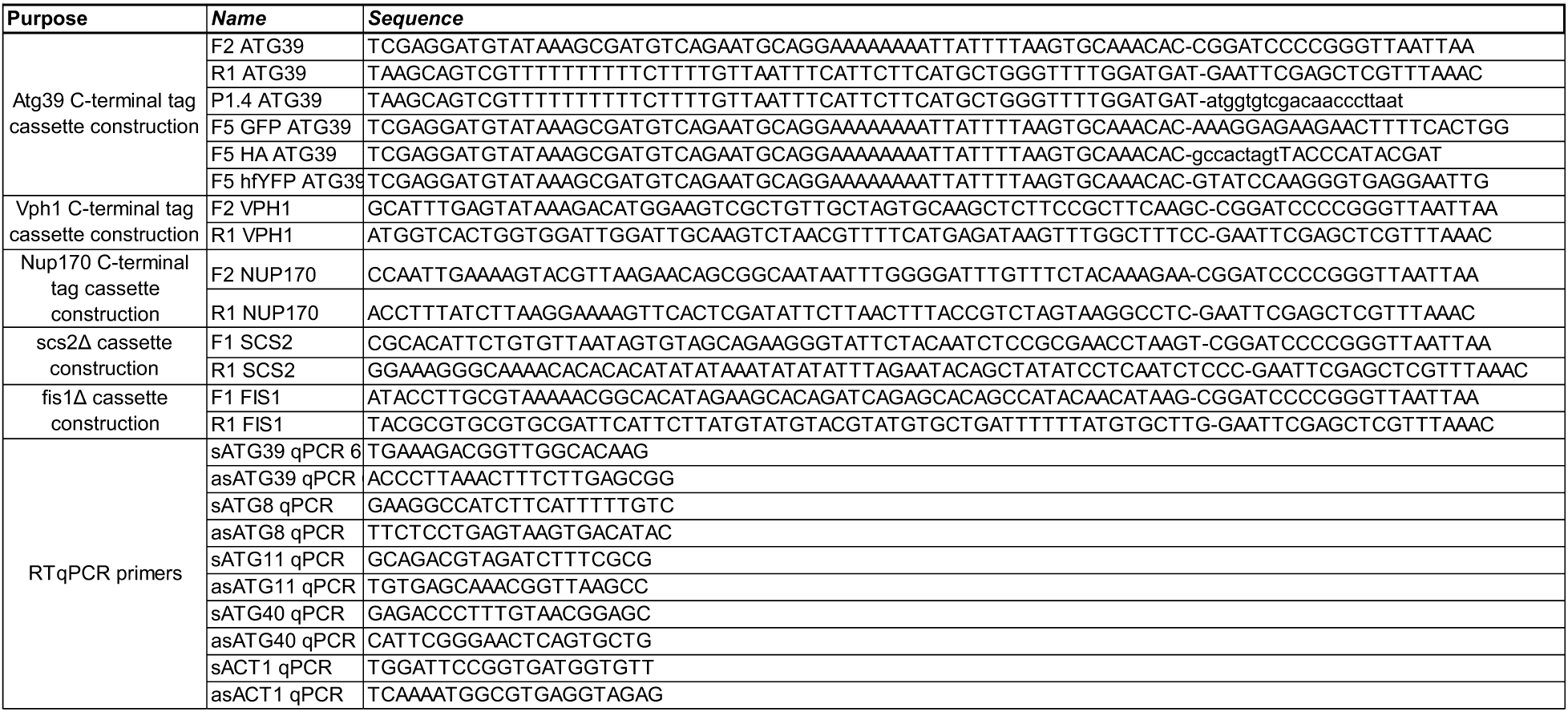
Primers used in this study.

## References

1. Mannino PJ, Lusk CP. Quality control mechanisms that protect nuclear envelope identity and function. J Cell Biol. Sep 5 2022;221(9)doi:10.1083/jcb.202205123

2. Mucino-Hernandez G, Acevo-Rodriguez PS, Cabrera-Benitez S, Guerrero AO, Merchant-Larios H, Castro-Obregon S. Nucleophagy contributes to genome stability through degradation of type II topoisomerases A and B and nucleolar components. J Cell Sci. Jan 1 2023;136(1)doi:10.1242/jcs.260563

3. Hasper J, Welle K, Swovick K, Hryhorenko J, Ghaemmaghami S, Buchwalter A. Long lifetime and tissue-specific accumulation of lamin A/C in Hutchinson-Gilford progeria syndrome. J Cell Biol. Jan 1 2024;223(1)doi:10.1083/jcb.202307049

4. Tsai PL, Zhao C, Turner E, Schlieker C. The Lamin B receptor is essential for cholesterol synthesis and perturbed by disease-causing mutations. Elife. Jun 23 2016;5doi:10.7554/eLife.16011

5. Papandreou ME, Tavernarakis N. Nucleophagy: from homeostasis to disease. Cell Death Differ. Mar 2019;26(4):630–639. doi:10.1038/s41418-018-0266-5

6. Natarajan N, Foresti O, Wendrich K, Stein A, Carvalho P. Quality Control of Protein Complex Assembly by a Transmembrane Recognition Factor. Mol Cell. Jan 2 2020;77(1):108–119 e9. doi:10.1016/j.molcel.2019.10.003

7. Khmelinskii A, Blaszczak E, Pantazopoulou M, et al. Protein quality control at the inner nuclear membrane. Nature. Dec 18 2014;516(7531):410–3. doi:10.1038/nature14096

8. Smoyer CJ, Smith SE, Gardner JM, McCroskey S, Unruh JR, Jaspersen SL. Distribution of Proteins at the Inner Nuclear Membrane Is Regulated by the Asi1 E3 Ligase in Saccharomyces cerevisiae. Genetics. Apr 2019;211(4):1269–1282. doi:10.1534/genetics.119.301911

9. Dou Z, Xu C, Donahue G, et al. Autophagy mediates degradation of nuclear lamina. Nature. Nov 5 2015;527(7576):105–9. doi:10.1038/nature15548

10. King GA, Goodman JS, Schick JG, et al. Meiotic cellular rejuvenation is coupled to nuclear remodeling in budding yeast. Elife. Aug 9 2019;8doi:10.7554/eLife.47156

11. King GA, Wettstein R, Varberg JM, et al. Meiotic nuclear pore complex remodeling provides key insights into nuclear basket organization. J Cell Biol. Feb 6 2023;222(2)doi:10.1083/jcb.202204039

12. Mizuno T, Irie K. Msn2/4 transcription factors positively regulate expression of Atg39 ER-phagy receptor. Sci Rep. Jun 7 2021;11(1):11919. doi:10.1038/s41598-021-91480-0

13. Papandreou ME, Konstantinidis G, Tavernarakis N. Nucleophagy delays aging and preserves germline immortality. Nat Aging. Jan 2023;3(1):34–46. doi:10.1038/s43587-022-00327-4

14. Wu N, Zheng W, Zhou Y, et al. Autophagy in aging-related diseases and cancer: Principles, regulatory mechanisms and therapeutic potential. Ageing Research Reviews. 2024;100doi:10.1016/j.arr.2024.102428

15. Bahmanyar S, Biggs R, Schuh AL, et al. Spatial control of phospholipid flux restricts endoplasmic reticulum sheet formation to allow nuclear envelope breakdown. Genes Dev. Jan 15 2014;28(2):121–6. doi:10.1101/gad.230599.113

16. Campbell JL, Lorenz A, Witkin KL, Hays T, Loidl J, Cohen-Fix O. Yeast nuclear envelope subdomains with distinct abilities to resist membrane expansion. Mol Biol Cell. Apr 2006;17(4):1768–78. doi:10.1091/mbc.e05-09-0839

17. Romanauska A, Kohler A. Lipid saturation controls nuclear envelope function. Nat Cell Biol. Sep 2023;25(9):1290–1302. doi:10.1038/s41556-023-01207-8

18. Santos-Rosa H, Leung J, Grimsey N, Peak-Chew S, Siniossoglou S. The yeast lipin Smp2 couples phospholipid biosynthesis to nuclear membrane growth. EMBO J. 2005;24

19. Barbosa AD, Lim K, Mari M, et al. Compartmentalized Synthesis of Triacylglycerol at the Inner Nuclear Membrane Regulates Nuclear Organization. Dev Cell. Sep 23 2019;50(6):755–766 e6. doi:10.1016/j.devcel.2019.07.009

20. Romanauska A, Kohler A. The Inner Nuclear Membrane Is a Metabolically Active Territory that Generates Nuclear Lipid Droplets. Cell. Jul 26 2018;174(3):700–715 e18. doi:10.1016/j.cell.2018.05.047

21. Romanauska A, Kohler A. Reprogrammed lipid metabolism protects inner nuclear membrane against unsaturated fat. Dev Cell. Sep 27 2021;56(18):2562–2578 e3. doi:10.1016/j.devcel.2021.07.018

22. Melia TJ, Lystad AH, Simonsen A. Autophagosome biogenesis: From membrane growth to closure. J Cell Biol. Jun 1 2020;219(6)doi:10.1083/jcb.202002085

23. Eickhorst C, Licheva M, Kraft C. Scaffold proteins in bulk and selective autophagy. Prog Mol Biol Transl Sci. 2020;172:15–35. doi:10.1016/bs.pmbts.2020.01.009

24. Kucinska MK, Fedry J, Galli C, et al. TMX4-driven LINC complex disassembly and asymmetric autophagy of the nuclear envelope upon acute ER stress. Nat Commun. Jun 13 2023;14(1):3497. doi:10.1038/s41467-023-39172-3

25. Roberts P, Moshitch-Moshkovitz S, Kvam E, O’Toole E, Winey M, Goldfarb DS. Piecemeal microautophagy of nucleus in Saccharomyces cerevisiae. Mol Biol Cell. Jan 2003;14(1):129–41. doi:10.1091/mbc.e02-08-0483

26. Allegretti M, Zimmerli CE, Rantos V, et al. In-cell architecture of the nuclear pore and snapshots of its turnover. Nature. Oct 2020;586(7831):796–800. doi:10.1038/s41586-020-2670-5

27. Lee CW, Wilfling F, Ronchi P, et al. Selective autophagy degrades nuclear pore complexes. Nat Cell Biol. Feb 2020;22(2):159–166. doi:10.1038/s41556-019-0459-2

28. Tomioka Y, Kotani T, Kirisako H, et al. TORC1 inactivation stimulates autophagy of nucleoporin and nuclear pore complexes. J Cell Biol. Jul 6 2020;219(7)doi:10.1083/jcb.201910063

29. Mochida K, Oikawa Y, Kimura Y, et al. Receptor-mediated selective autophagy degrades the endoplasmic reticulum and the nucleus. Nature. Jun 18 2015;522(7556):359–62. doi:10.1038/nature14506

30. Mizuno T, Muroi K, Irie K. Snf1 AMPK positively regulates ER-phagy via expression control of Atg39 autophagy receptor in yeast ER stress response. PLoS Genet. Sep 2020;16(9):e1009053. doi:10.1371/journal.pgen.1009053

31. Otto FB, Thumm M. Mechanistic dissection of macro- and micronucleophagy. Autophagy. Mar 2021;17(3):626–639. doi:10.1080/15548627.2020.1725402

32. Chandra S, Mannino PJ, Thaller DJ, et al. Atg39 selectively captures inner nuclear membrane into lumenal vesicles for delivery to the autophagosome. J Cell Biol. Dec 6 2021;220(12)doi:10.1083/jcb.202103030

33. Mochida K, Otani T, Katsumata Y, et al. Atg39 links and deforms the outer and inner nuclear membranes in selective autophagy of the nucleus. J Cell Biol. Feb 7 2022;221(2)doi:10.1083/jcb.202103178

34. Otsuga D, Keegan BR, Brisch E, et al. The Dynamin-related GTPase, Dnm1p, Controls Mitochondrial Morphology in Yeast. J Cell Biol. 1998;143(2)

35. Kuravi K, Nagotu S, Krikken AM, et al. Dynamin-related proteins Vps1p and Dnm1p control peroxisome abundance in Saccharomyces cerevisiae. J Cell Sci. Oct 1 2006;119(Pt 19):3994–4001. doi:10.1242/jcs.03166

36. Gonzalez A, Covarrubias-Pinto A, Bhaskara RM, et al. Ubiquitination regulates ER-phagy and remodelling of endoplasmic reticulum. Nature. Jun 2023;618(7964):394–401. doi:10.1038/s41586-023-06089-2

37. Rosado CJ, Mijaljica D, Hatzinisiriou I, Prescott M, Devenish RJ. Rosella: a fluorescent pH-biosensor for reporting vacuolar turnover of cytosol and organelles in yeast. Autophagy. Feb 2008;4(2):205–13. doi:10.4161/auto.5331

38. Kim J, Kamada Y, Stromhaug PE, et al. Cvt9/Gsa9 Functions in Sequestering Selective Cytosolic Cargo Destined for the Vacuole. Journal of Cell Biology. 2001;153(1)

39. Kirisako T, Baba M, Ishihara N, et al. Formation Process of Autophogosome is Traced with Apg8/Aut7p in Yeast. Journal of Cell Biology. 1999;147(2)

40. Kvam E, Goldfarb DS. Nvj1p is the outer-nuclear-membrane receptor for oxysterol-binding protein homolog Osh1p in Saccharomyces cerevisiae. J Cell Sci. Oct 1 2004;117(Pt 21):4959–68. doi:10.1242/jcs.01372

41. Borah S, Thaller DJ, Hakhverdyan Z, et al. Heh2/Man1 may be an evolutionarily conserved sensor of NPC assembly state. Mol Biol Cell. Jul 15 2021;32(15):1359–1373. doi:10.1091/mbc.E20-09-0584

42. Thaller DJ, Allegretti M, Borah S, Ronchi P, Beck M, Lusk CP. An ESCRT-LEM protein surveillance system is poised to directly monitor the nuclear envelope and nuclear transport system. Elife. Apr 3 2019;8doi:10.7554/eLife.45284

43. Webster BM, Colombi P, Jager J, Lusk CP. Surveillance of nuclear pore complex assembly by ESCRT-III/Vps4. Cell. Oct 9 2014;159(2):388-401. doi:10.1016/j.cell.2014.09.012

44. Webster BM, Thaller DJ, Jager J, Ochmann SE, Borah S, Lusk CP. Chm7 and Heh1 collaborate to link nuclear pore complex quality control with nuclear envelope sealing. EMBO J. 2016;35

45. Chen S, Cui Y, Parashar S, Novick PJ, Ferro-Novick S. ER-phagy requires Lnp1, a protein that stabilizes rearrangements of the ER network. Proc Natl Acad Sci U S A. Jul 3 2018;115(27):E6237–E6244. doi:10.1073/pnas.1805032115

46. Chen S, Mari M, Parashar S, et al. Vps13 is required for the packaging of the ER into autophagosomes during ER-phagy. Proc Natl Acad Sci U S A. Aug 4 2020;117(31):18530–18539. doi:10.1073/pnas.2008923117

47. Cui Y, Parashar S, Zahoor M, et al. A COPII subunit acts with an autophagy receptor to target endoplasmic reticulum for degradation. Science. Jul 5 2019;365(6448):53–60. doi:10.1126/science.aau9263

48. Liu D, Mari M, Li X, Reggiori F, Ferro-Novick S, Novick P. ER-phagy requires the assembly of actin at sites of contact between the cortical ER and endocytic pits. Proc Natl Acad Sci U S A. Feb 8 2022;119(6)doi:10.1073/pnas.2117554119

49. Anwar K, Klemm RW, Condon A, et al. The dynamin-like GTPase Sey1p mediates homotypic ER fusion in S. cerevisiae. J Cell Biol. Apr 16 2012;197(2):209–17. doi:10.1083/jcb.201111115

50. Voeltz GK, Prinz WA, Shibata Y, Rist JM, Rapoport TA. A class of membrane proteins shaping the tubular endoplasmic reticulum. Cell. Feb 10 2006;124(3):573–86. doi:10.1016/j.cell.2005.11.047

51. Mozdy AD, McCaffery JM, Shaw JM. Dnm1p GTPase-mediated Mitochondrial Fission Is a Multi-step Process Requiring the Novel Integral Membrane Component Fis1p. J Cell Biol. 2000;151

52. Sesaki H, Southard SM, Yaffe MP, Jensen RE. Mgm1p, a dynamin-related GTPase, is essential for fusion of the mitochondrial outer membrane. Mol Biol Cell. Jun 2003;14(6):2342–56. doi:10.1091/mbc.e02-12-0788

53. Smaczynska-de R, II, Allwood EG, Aghamohammadzadeh S, Hettema EH, Goldberg MW, Ayscough KR. A role for the dynamin-like protein Vps1 during endocytosis in yeast. J Cell Sci. Oct 15 2010;123(Pt 20):3496–506. doi:10.1242/jcs.070508

54. Ferguson SM, De Camilli P. Dynamin, a membrane-remodelling GTPase. Nat Rev Mol Cell Biol. Jan 11 2012;13(2):75–88. doi:10.1038/nrm3266

55. Mao K, Liu X, Feng Y, Klionsky DJ. The progression of peroxisomal degradation through autophagy requires peroxisomal division. Autophagy. Apr 2014;10(4):652–61. doi:10.4161/auto.27852

56. Motley AM, Ward GP, Hettema EH. Dnm1p-dependent peroxisome fission requires Caf4p, Mdv1p and Fis1p. J Cell Sci. May 15 2008;121(Pt 10):1633–40. doi:10.1242/jcs.026344

57. Naylor K, Ingerman E, Okreglak V, Marino M, Hinshaw JE, Nunnari J. Mdv1 Interacts with Assembled Dnm1 to Promote Mitochondrial Division. Journal of Biological Chemistry. 2006;281(4):2177–2183. doi:10.1074/jbc.M507943200

58. Mao K, Wang K, Liu X, Klionsky DJ. The scaffold protein Atg11 recruits fission machinery to drive selective mitochondria degradation by autophagy. Dev Cell. Jul 15 2013;26(1):9–18. doi:10.1016/j.devcel.2013.05.024

59. Hu C, Chinenov Y, Kerppola TK. Visualization of Interactions among bZIP and Rel Family Proteins in Living Cells Using Bimolecular Fluorescence Complementation. Molecular Cell. 2002;9

60. Ader NR, Chen L, Surovtsev IV, et al. An ESCRT grommet cooperates with a diffusion barrier to maintain nuclear integrity. Nat Cell Biol. Oct 2023;25(10):1465–1477. doi:10.1038/s41556-023-01235-4

61. Bailey MLP, Surovtsev I, Williams JF, et al. Loops and the activity of loop extrusion factors constrain chromatin dynamics. Mol Biol Cell. Jul 1 2023;34(8):ar78. doi:10.1091/mbc.E23-04-0119

62. Crocker JC, Grier DG. Methods of digital video microscopy for colloidal studies. Journal of colloid and interface science. 1996;179

63. Lawrimore J, Bloom KS, Salmon ED. Point centromeres contain more than a single centromere-specific Cse4 (CENP-A) nucleosome. J Cell Biol. Nov 14 2011;195(4):573–82. doi:10.1083/jcb.201106036

64. Campbell BC, Paez-Segala MG, Looger LL, Petsko GA, Liu CF. Chemically stable fluorescent proteins for advanced microscopy. Nat Methods. Dec 2022;19(12):1612–1621. doi:10.1038/s41592-022-01660-7

65. Friedman JR, Lackner LL, West M, R. DJ, Nunnari J, Voeltz GK. ER Tubules Mark Sites of Mitochondrial Division. Science. 2011;334

66. Villinger C, Neusser G, Kranz C, Walther P, Mertens T. 3D Analysis of HCMV Induced-Nuclear Membrane Structures by FIB/SEM Tomography: Insight into an Unprecedented Membrane Morphology. Viruses. Nov 4 2015;7(11):5686–704. doi:10.3390/v7112900

67. Speese SD, Ashley J, Jokhi V, et al. Nuclear envelope budding enables large ribonucleoprotein particle export during synaptic Wnt signaling. Cell. May 11 2012;149(4):832–46. doi:10.1016/j.cell.2012.03.032

68. Fujioka Y, Tsuji T, Kotani T, et al. 2024;doi:10.1101/2024.08.29.610189

69. Schneider BL, Seufert W, Steiner B, Yang QH, Futcher AB. Use of polymerase chain reaction epitope tagging for protein tagging in Saccharomyces cerevisiae. Yeast. Oct 1995;11(13):1265–74. doi:10.1002/yea.320111306

70. Longtine MS, McKenzie Iii A, Demarini DJ, et al. Additional modules for versatile and economical PCR-based gene deletion and modification in Saccharomyces cerevisiae. Yeast. 1998;14(10):953–961. doi:10.1002/(sici)1097-0061(199807)14:10<953::Aid-yea293>3.0.Co;2-u

71. Zhang Y, Serratore ND, Briggs SD. N-ICE plasmids for generating N-terminal 3 x FLAG tagged genes that allow inducible, constitutive or endogenous expression in Saccharomyces cerevisiae. Yeast. May 2017;34(5):223–235. doi:10.1002/yea.3226

72. Baudin A, Ozier-Kalogeropoulos OD, A., Lacroute F, Cullin C. A simple and efficient method for direct gene deletion in *Saccharomyces cerevisiae*. Nucleic Acids Research. 1993;21

73. Guldener U, Heck S, Fiedler T, Beinhauer J, Hegemann JH. A new efficient gene disruption cassette for repeated use in budding yeast. Nucleic Acids Research. 1996;24(13)

74. Lacy MM, Baddeley D, Berro J. Single-molecule imaging of the BAR-domain protein Pil1p reveals filament-end dynamics. Mol Biol Cell. Aug 15 2017;28(17):2251–2259. doi:10.1091/mbc.E17-04-0238

75. Smoyer CJ, Katta SS, Gardner JM, et al. Analysis of membrane proteins localizing to the inner nuclear envelope in living cells. J Cell Biol. Nov 21 2016;215(4):575–590. doi:10.1083/jcb.201607043

76. Chen BC, Legant WR, Wang K, et al. Lattice light-sheet microscopy: imaging molecules to embryos at high spatiotemporal resolution. Science. Oct 24 2014;346(6208):1257998. doi:10.1126/science.1257998

77. Adell MAY, Migliano SM, Upadhyayula S, et al. Recruitment dynamics of ESCRT-III and Vps4 to endosomes and implications for reverse membrane budding. Elife. Oct 11 2017;6doi:10.7554/eLife.31652

78. Kukulski W, Schorb M, Welsch S, Picco A, Kaksonen M, Briggs JA. Precise, correlated fluorescence microscopy and electron tomography of lowicryl sections using fluorescent fiducial markers. Methods Cell Biol. 2012;111:235–57. doi:10.1016/B978-0-12-416026-2.00013-3

79. Mastronarde DN. Automated electron microscope tomography using robust prediction of specimen movements. J Struct Biol. Oct 2005;152(1):36–51. doi:10.1016/j.jsb.2005.07.007

80. Kremer JRM, D. N. McIntosh J. R. Computer Visualization of Three-Dimensional Image Data Using IMOD. Journal of Structural Biology. 1996;116(1)

81. Mastronarde DN, Held SR. Automated tilt series alignment and tomographic reconstruction in IMOD. J Struct Biol. Feb 2017;197(2):102–113. doi:10.1016/j.jsb.2016.07.011

82. Flajolet P, Gardy D, Thimonier L. Birthday paradox, coupon collectors, caching algorithms and self-organizing search. Discrete Applied Mathematics. 1992;39:207–229.

83. Schindelin J, Arganda-Carreras I, Frise E, et al. Fiji: an open-source platform for biological-image analysis. Nat Methods. Jun 28 2012;9(7):676–82. doi:10.1038/nmeth.2019

84. Bolte S, Cordelieres FP. A guided tour into subcellular colocalization analysis in light microscopy. Journal of Microscopy. 2006;224doi:10.1111/j.1365-2818.2006.01706.x

